# Maturation of Gait: Identification of Locomotor Profiles from Early Childhood to Adulthood

**DOI:** 10.64898/2026.06.02.729581

**Authors:** Silvère De Freitas, Mohammad Reza Effatparvar, Fabien Dal Maso, Yosra Cherni

## Abstract

**Background:** Human gait is a key marker of motor development. While walking on even surface is well-documented, responses to irregular surfaces, closer to real-world environments, remain understudied. This limitation is reinforced by the frequent use of univariate analyses, though locomotor control emerges from interactions of multiple features. Multivariate approaches are therefore essential to characterize developmental modulations to uneven surfaces.

**Objectives:** (i) To evaluate the combined effects of age and surface complexity on multiple gait domains in healthy individuals during development; (ii) To identify locomotor profiles across gait maturation.

**Methods:** Sixty-eight participants (2-35 years) walked at a self-selected speed on even, medium, and high irregularity surfaces. Gait kinematics were captured using a 3D motion system. Linear Mixed Models evaluated age and surface effects on 28 variables across five domains: pace, rhythm, dynamic stability, variability, and asymmetry. Moreover, principal component analysis followed by *k*-means clustering was performed on 15 normalized variables to identify gait profiles.

**Results:** Age and surface influenced most variables (*p*<0.05). Young children (2-5 years) exhibited the greatest modulation of asymmetry, base of support, smoothness, and dynamic stability with surface complexity. Conversely, adults and adolescents (12-35 years) showed higher variability modulation on irregular surface. PCA-assisted clustering identified two clusters: Cluster1 (15.2 years, smooth-regular) and Cluster2 (6.1 years, wide-base-variable). Across surfaces, five subgroups emerged: two *consistent* (15.8 years in Cluster1; 5.4 years in Cluster2) and three *switchers* (8.5 years [7.6-12.8]) showing context-dependent transitions as surface complexity increased.

**Discussion:** Differential maturation and surface sensitivity suggest that irregular surfaces act as functional stressors, revealing developmental gaps hidden on even ground. The surface-dependent transition at 7-13 years suggests that locomotor maturity is task-dependent rather than a fixed state, shifting from stable, regular to a variability-driven, balance-supportive strategy with complexity. These profiles delineate developmental stages and may help to identify atypical trajectories in pediatric rehabilitation.

## 1. Introduction

Locomotor maturation follows a well-established developmental trajectory. Independent walking typically emerges around the end of the first year of life (1). Within the first months of independent walking, the temporal organization of gait become progressively organized, as reflected by the convergence of stride/stance, stance/swing and swing/double support duration ratios toward adult-like values (2). During this period, gait maturation is influenced by walking experience as well as chronological age, and is accompanied by an increase of single-limb stance duration (2,3). This fundamental rhythmic structure of locomotion is followed by a progressive increase in gait speed and step length, together with a reduction in cadence, step width, and the time spent in double support, reflecting a transition from a cautious gait toward a more efficient pattern of forward progression during the first 5 years of childhood (3–6). Although gait speed has been reported to approach adult-like values around 7 years of age (3,7), gait maturation continues beyond this period, notably with a progressive reduction in step-to-step variability observed throughout late childhood and adolescence while variables such as cadence, step length, and support times continue to evolve in relation to ongoing lower-limb growth and progressive adjustments in locomotor control (8). Adolescents up to 17 years may still differ from adults, as reflected by the Gait Variability Index (GVI), a composite measure derived from nine spatiotemporal variables (9). In parallel, gait organization may continue to be refined during adolescence, particularly during periods of rapid growth, as reflected by changes in trunk-acceleration-derived measures of gait smoothness and step regularity (10,11). Yet, the majority of studies focusing on gait maturation have been conducted under steady conditions on even surfaces which may lead to the interpretation that some gait features are mature although their robustness under more demanding environmental constraints remains untested. As a result, how locomotor control matures in ecological environments that require continuous sensorimotor adjustments remains poorly understood (12). This issue may be further compounded by age-based classifications, which can mask inter-individual variability in adaptive locomotor strategies during development.

Beyond age-related maturation, locomotor pattern must be continuously adjusted to external constraints that challenge balance and coordination. When walking on uneven surface, individuals must adjust foot placement, regulate step timing, and modulate body dynamics to maintain balance while generating forward propulsion (1). Previous studies have consistently shown that uneven surfaces decrease walking speed, step length, and cadence, and increase step width to maintain dynamic stability in mechanically unpredictable environments (13–15). In addition to these spatiotemporal adjustments, walking on uneven surface has been associated with increased stride-to-stride variability (16–18) and reduced gait smoothness (19,20), indicating the need for continuous sensorimotor adjustments from one step to the next. While such variability is widely recognized as a key feature of adaptive locomotor control, as it enables flexible responses to environmental constraints rather than simply reflecting motor noise (18), changes in gait smoothness may reflect modifications in the quality and temporal regulation of movement, as smoothness is sensitive to the continuity and corrective processes of locomotor control (21). Together, these gait responses suggest that the ability to adjust gait to irregular surfaces is an important component of locomotor control. Therefore, gait maturation cannot be fully understood without considering how individuals respond to mechanically challenging environments.

Most previous studies have examined these locomotor responses using univariate or domain-specific approaches, implicitly assuming that gait variables evolve independently. Furthermore, participants are typically grouped into discrete age bands (e.g., < 5, 6-11, < 18y) rather than being analyzed as a continuum, which may mask meaningful within-group variability (11,22). Such approaches may overlook the coordinated organization of locomotor control, where multiple domains interact to produce stable and efficient movement patterns. As a result, two individuals may walk at a similar speed while using different combinations of step length, cadence, step width, variability, or stability-related adjustments. An isolated variable may therefore suggest similar performance, whereas the overall gait organization may differ substantially. Addressing this gap requires analyses capable of capturing the multidimensional nature of gait adjustments, involving multiple domains such as pace, rhythm, dynamic stability, variability, and asymmetry (12). This complexity, combined with the large volume of data generated by quantitative gait analysis, complicates interpretation and limits clinical uptake. Multivariate approaches, such as principal component analysis (PCA) and clustering algorithms, may provide a means to synthesize high-dimensional data and identify locomotor profiles, which can be defined as patterns emerging from coordinated organization of gait domains rather than from chronological age alone (23,24). Such profiling approaches have been applied across various locomotor tasks and linked to functional outcomes (23,25,26). In the context of gait maturation, they may help reveal how multiple gait dimensions are organized into developmental locomotor strategies.

The present study aimed to identify profiles of gait maturation while walking on different types of surfaces. To this end, we defined two complementary objectives, each addressed through a distinct analytical approach. Our first objective was to evaluate the combined effects of age and walking surface complexity on gait pace, rhythm, dynamic stability, variability, and asymmetry in healthy individuals using a hypothesis-driven analysis. We hypothesized that increasing surface complexity would induce modifications across multiple domains of gait including variability, stability, and rhythm-related smoothness, and these modifications would vary across developmental stages. The second objective was to identify distinct locomotor profiles across gait maturation using an exploratory data-driven analysis. We hypothesized that multivariate analyses would reveal distinct locomotor profiles primarily driven by maturation and that increasing surface complexity would modulate profile expression by revealing context-dependent locomotor adjustments.

## 2. Materials and Methods

### 2.1. Participants

Sixty-eight healthy volunteers aged between 2 and 35 years were recruited from the local community to participate to this study. This lower age limit was chosen for safety and feasibility reasons, as the protocol required participants to walk without assistance across irregular surfaces. The upper age limit was fixed at 35 years consistent with classification of young adulthood in gait literature (27,28). Participants unable to understand simple instructions, with musculoskeletal diseases or previous orthopedic surgery, or a medical or neurologic disease affecting age-related normal gait were excluded. All experimental procedures were approved by the Research Ethics Board of Sainte-Justine Hospital (2026-9359). For minors, parents provided written informed consent, and children provided verbal assent which was recorded in the study documentation by the investigator before participation. Adult participants provided written informed consent (29). Participants were recruited between December 4, 2024, and December 12, 2025.

### 2.2. Data Collection

#### Kinematics

Whole-body kinematics were captured at 100 Hz using a 12-camera optoelectronic motion capture system (Vicon Motion Systems Ltd., Oxford, UK). Forty-three reflective markers were placed on anatomical landmarks according to the Plug-in Gait Full Body model. To scale the biomechanical model, body mass, standing height, and leg length (L0) were measured to the nearest 0.1 kg and 0.1 cm.

#### Experimental conditions

Participants walked at a comfortable self-selected speed across a 4.2 × 1 m walkway under three surface conditions (Fig. 1): an **Even, Medium,** and **High**. The Even condition consisted in walking on the laboratory floor. The Medium and High conditions consisted of walking on shock-absorbing polyurethane panels with their introduction ramps (Terrasensa, Kassel, Germany). The panels had a Shore A hardness of 45 on a 0-100 scale, indicating a moderately compliant semi-rigid polyurethane material. Maximum vertical variation of 2 cm and 5 cm created surface irregularity for the Medium and High surfaces, respectively. To exclude acceleration and deceleration gait phases from subsequent analysis, a 2 m flat walkway was provided before and after the 4.2 m even, medium, and high walkways. Children, adolescents, and adults performed 10 trials. Young children performed five trials to limit fatigue and ensure compliance. This did not affect the analysis, as young children’s shorter height resulted in more gait cycles per trial than in older groups. Surfaces were presented in a randomized order between participants. All participants wore their own athletic footwear during testing.

**Figure 1:**
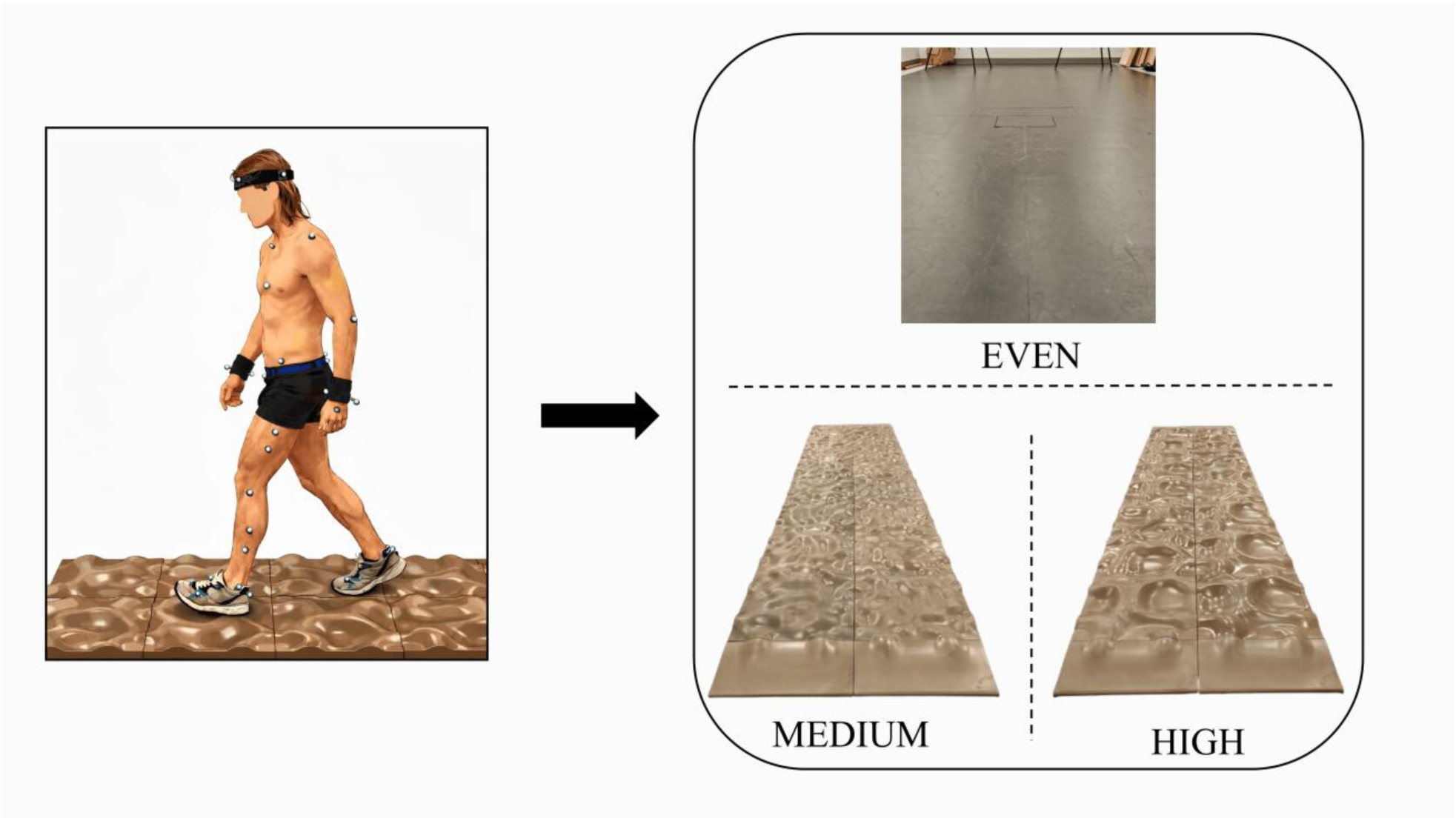
Experimental setup and surface conditions. Left: Participant equipped with the full-body reflective marker set (Plug-in Gait model) walking on the uneven surface panels. Top right: The Even surface of the laboratory floor. Bottom right: Medium and High polyurethane panels.

### 2.3. Data processing

Reflective marker labeling and gap filling were performed using Vicon Nexus software (v2.14.0, Vicon Motion Systems Ltd., Oxford, UK). Foot-strike and foot-off events were manually identified by a single experimented investigator for all participants and conditions. Following event detection, marker trajectories data segmented into individual gait cycles. To ensure a balanced statistical design, the number of gait cycles retained for analysis was set to the minimum number of cycles available across legs and surfaces for each participant. Each participant had at least 10 recorded cycles per condition prior to this equalization. This threshold was selected to exceed the minimum of eight cycles recommended to achieve good reliability for most spatiotemporal and stability variables investigated in the present study (30). Thus, statistical analyses were based on 23 ± 7 gait cycles per surface in young children (left leg: 11 ± 3; right leg: 12 ± 3), 33 ± 9 gait cycles in children (left leg: 17 ± 4; right leg: 16 ± 5), 23 ± 5 gait cycles in adolescents (left leg: 12 ± 2; right leg: 11 ± 3), and 24 ± 5 gait cycles in adults (left leg: 12 ± 3; right leg: 12 ± 3).

### 2.4. Variables’ calculation

A total of 28 gait variables were calculated and categorized into 5 domains, namely: Pace (7 variables), Rhythm (9 variables), Dynamic stability (6 variables), Variability (3 variables), and Asymmetry (3 variables) (31). For all subsequent analyses, variables with separate left and right limb values were averaged across limbs. To account for differences in body size between participants, variables were normalized to lower limb length according to Hof’s dynamic similarity principles (32). All normalized variables are dimensionless.

#### Pace domain

This domain refers to forward progression. It includes:

- raw (m/s) and normalized gait speed (au). Gait speed was calculated from the antero-posterior (AP) distance traveled by the center of mass (CoM) estimated as the centroid of the 4 reflective markers positioned on the pelvis;
- stride and step length (m);
- normalized step length (au);
- raw (cm.min.step^-1^) and normalized (au) walk ratio. Walk ratio was calculated as the ratio between step length and cadence.

#### Rhythm domain

This domain refers to the temporal organization, periodicity, and smoothness of gait. It includes:

- double and single support time (expressed as a % of gait cycle);
- stance time (s);
- swing time (s);
- step and stride time (s);
- raw (steps.min^-1^) and normalized cadence (au);
- the spectral arc length (SPARC) index was used to quantify gait smoothness from the frequency content of the CoM velocity profile (21). For each trial, SPARC was calculated over the walking segment delimited by the first and last heel strikes. The three-dimensional CoM velocity magnitude was demeaned, transformed into the frequency domain using a fast Fourier transform with zero-padded using an index of 4, and normalized by its maximum spectral amplitude within the 0-6 Hz band. SPARC was then calculated as the negative arc length of the normalized power spectrum magnitude, using an amplitude threshold of 0.05 (to retain only the portion of the spectrum containing meaningful movement-related content and limiting noise) and a maximum frequency of 6 Hz (21). Less negative SPARC values indicate smoother gait.

#### Dynamic stability domain

This domain refers to the ability to maintain dynamic stability during forward progression. It includes:

- raw (cm) and normalized step width;
- raw (mm) and normalized margin of stability (MoS) in both AP and medio-lateral (ML) directions at heel strike. The MoS was calculated as the distance between the extrapolated CoM and the boundaries of the base of support (33–36). For AP MoS, the anterior boundary of the MoS was defined using the heel marker, whereas for ML MoS, the lateral boundary was defined using the lateral ankle marker (37).

#### Variability domain

This domain refers to the step-to-step variability. It includes:

- Gait variability index (GVI, (38)); GVI was calculated as a composite measure of spatiotemporal gait variability including nine variables: step length, stride length, step time, stride time, swing time, stance time, single support time, double support time, and stride velocity. For each limb, variables were normalized to their mean value, and variability was derived from the mean and standard deviation of the absolute differences between consecutive gait cycles, weighted according to Gouelle et al. (38). Left and right limb scores were averaged before computing the final GVI. In the present study, the adult group walking on the Even surface was used as the reference population to normalize all GVI values so that a GVI of 100 corresponds to the reference mean, whereas lower values indicate greater gait variability.
- coefficient of variation (CV, %) for step width and gait speed.

#### Asymmetry domain

This domain refers to functional differences between the left and right lower limbs. It includes:

- Robinson’s Symmetry Index (SI, %) for double support time, stride length, and step width (39), as follow (Equation 1):

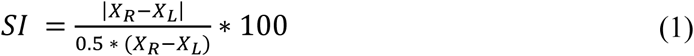

where 𝑋_𝑅_ and 𝑋_𝐿_ represent the values of the gait variable for the right and left limbs, respectively.

### 2.5. Statistical analysis

#### 2.5.1. Hypothesis-driven analysis

Statistical analyses were performed using R (version 4.5.2). The level of significance was set at α = 0.05 for all tests. Magnitude of the effects was evaluated using partial eta-squared (η²p), where values of 0.01, 0.06, and 0.14 were considered small, medium, and large effects, respectively (40) or *d* values where <0.20, 0.20-0.49, 0.5-0.79, and >0.8 were considered trivial, small, medium, and large effects, respectively, where appropriate (40).

##### Participants’ demography

One-way analyses of variance (ANOVA) were performed to assess differences in age and anthropometric characteristics (height, body mass, and L0) across age groups. Participants were allocated *a priori* into four groups (41): Young Children (2-5 years), Children (6-11 years), Adolescents (12-17 years), and Adults (18-35 years). These categories broadly correspond to early childhood, middle childhood, adolescence, and young adulthood, respectively, and were selected to capture major periods of growth and developmental change (42). Sex distribution was compared across age groups using Fisher’s exact test (43).

##### Gait variables

A linear mixed-effects model (LMM) was used to assess the effects of Age, Surface, and their interaction on each gait variable. Models were fitted using the *lme4* package (R), with gait variables entered as dependent variables, Age and Surface and their interaction as fixed effects, and Participant as a random intercept. When significant main effects or interactions were detected, post-hoc pairwise comparisons were performed using estimated marginal means, with Holm-Bonferroni correction. P-values were additionally adjusted using a False Discovery Rate (FDR) procedure (q = 0.05, FDR = 5%) (44). For post-hoc pairwise comparisons, effect sizes were reported using Cohen’s *d*, calculated with the model’s residual standard deviation as the denominator. Additionally, global model variance was assessed using Nakagawa’s marginal and conditional R² (R²*_m_* and R²*_c_*, respectively).

#### 2.5.2. Data-driven analysis: Locomotor profiles identification

To reduce dimensionality and redundancy among correlated gait variables, a PCA was performed on the normalized dataset using a singular value decomposition algorithm (45). Prior, features analyzed were selected *a priori* according to the two following methodological criteria. First, only normalized variables were selected in order to reduce anthropometric effects considering that height varies considerably between young children and adults. This reduced the initial set of 28 to 16 variables across the five domains: pace (n=3 variables), rhythm (n=4 variables), dynamic stability (n=3 variables), variability (n=3 variables), and asymmetry (n=3 variables). Second, to avoid overrepresentation of the rhythm domain that included 4 normalized variables, we investigated the correlations between all pairs of rhythm-related variables. Double and single support time showed the greatest correlation (r = 0.99 for all surfaces, while r < 0.86 for all other correlations). We removed single support time. The 15 gait variables retained for the PCA-assisted clustering analysis were:

- Pace domain: normalized gait speed, step length, and walk ratio;
- Rhythm domain: double support time, normalized cadence, and SPARC;
- Dynamic stability domain: normalized step width and AP and ML MoS;
- Variability domain: GVI, CV of step width, and gait speed;
- Asymmetry domain: SI of stride length, double support time, and step width.

To establish a global locomotor space, data from all age groups (Young children, Children, Adolescents, Adults) and walking surfaces (Even, Medium, High) were pooled. Before analysis, all variables were z-scored to ensure that the outcome of PCA-assisted clustering was driven by the relative variance of each gait variable rather than their absolute magnitudes (45,46). A sufficient number of principal components (PCs) were retained to explain at least 70% of the total variance (47). Then, the *k*-means clustering algorithm was applied to the retained PC scores with 50 replicates (independent runs from different random initializations, retaining the solution minimizing the within-cluster sum of squares), 500 maximum iterations (convergence limit within each run), and a fixed random seed for reproducibility. Cluster stability was assessed using a 100-iteration bootstrap procedure. To determine the optimal number of clusters (*k*), we evaluated the inertia curve (elbow method) (48,49) on a range of k = 1 to 10, with k = 1 used as the baseline for assessing the marginal decrease in within-cluster sum of squares. Further, the silhouette score (50), and Adjusted Rand Index (ARI) (51) were evaluated on a range of k = 2 to 10, since these metrics are only meaningful for partitions with at least two clusters.

##### Clustering and Pattern Recognition

Differences in demographic distribution (Age, Sex) and experimental conditions (Surface) between clusters were explored as well as changes in participants’ assignment into clusters across Even, Medium, High surfaces. When heterogeneity in transition patterns was observed between surfaces, exploratory subgroup analyses were conducted to further characterize transition profiles.

## 3. Results

### 3.1. Participants

Participant demographics are summarized in Table 1. One-way ANOVAs revealed a significant and large main effect of Age on age and all anthropometric characteristics: age, height, body mass, and leg length (L0) (all p < 0.001 and η²p > 0.50). Pairwise post-hoc analyses revealed that height and L0 differed significantly across Age, except between adolescents and adults (height: p = 0.15; L0: p = 0.50). Body mass differed significantly across age (p < 0.05). No significant association was observed between sex and Age (Fisher’s exact test, p = 0.79).

**Table 1:**
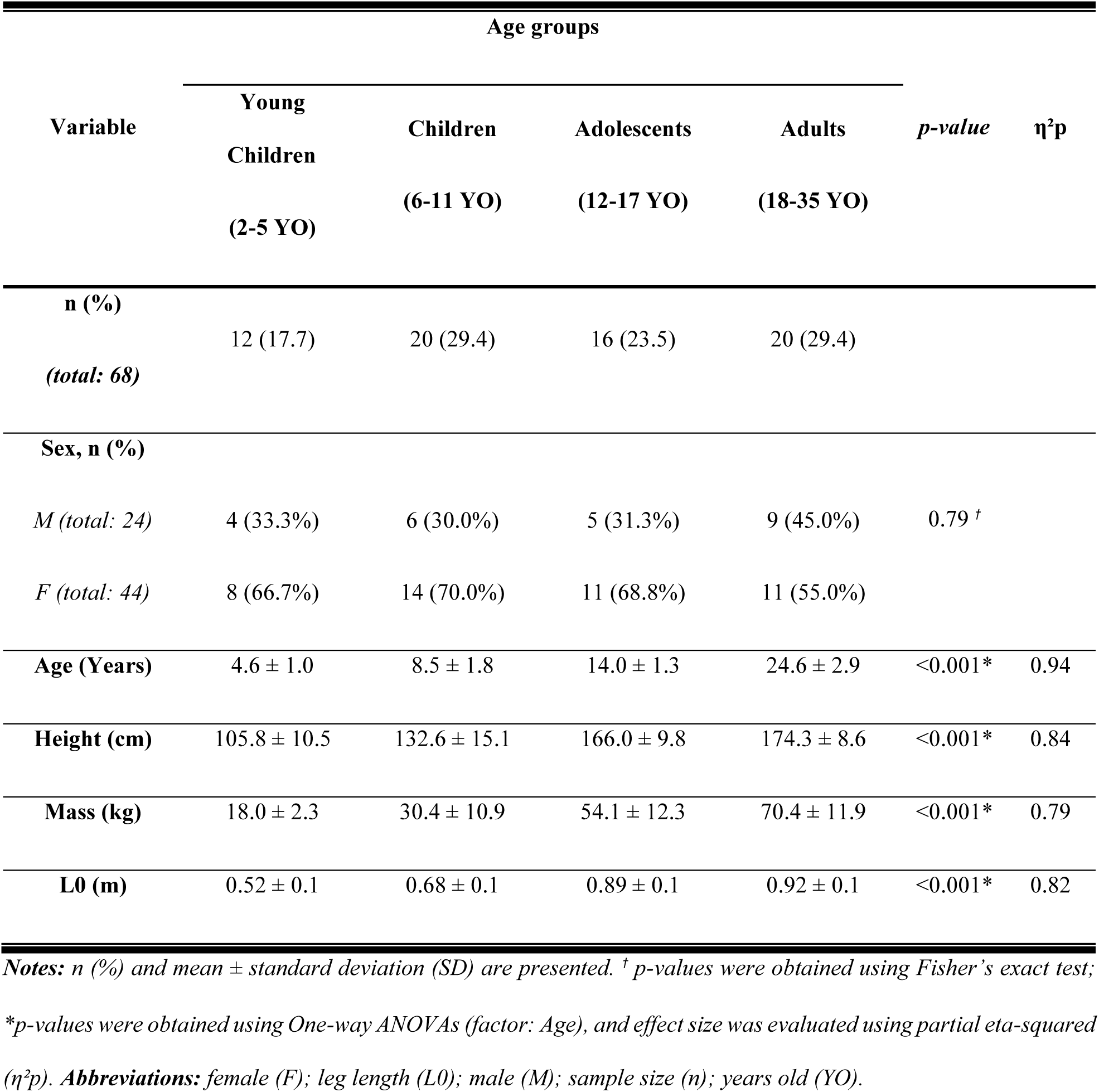
Demographic characteristics across age groups.

### 3.2. Gait variables

LMM revealed significant main effects of Age and Surface conditions across multiple gait variables, whereas Age × Surface interactions were limited to a subset of variables as presented in Table S2, which presents the descriptive values of variables per gait domain according to age group and surface condition. Effect size (*η²p, R²_m_,* and *R²_c_*) for each variable was further described in the Table S3. Post-hoc pairwise comparisons between age groups by Surface, and between Surface by age groups for gait variables showing a significant effect of the interaction Age × Surface are presented in Table S4 and S5 respectively. Table S6 reported post-hoc pairwise comparisons between each age group and adults for gait variables showing a significant effect of Age, and Table S7 post-hoc pairwise comparisons between each irregular surface (Medium, High) and even for gait variables showing a significant effect of Surface.

#### Pace domain

There was a significant Age × Surface interaction for normalized step length (q = 0.02, η²p = 0.13). Normalized step length significantly decreased with increasing surface irregularity only in young children (p < 0.001, d = 1.10-2.32). On the even surface, young children and children showed significantly higher normalized step length than adolescents and adults (p < 0.001, *d* = 3.28 and 3.23, respectively (Fig. 2).

**Figure 2:**
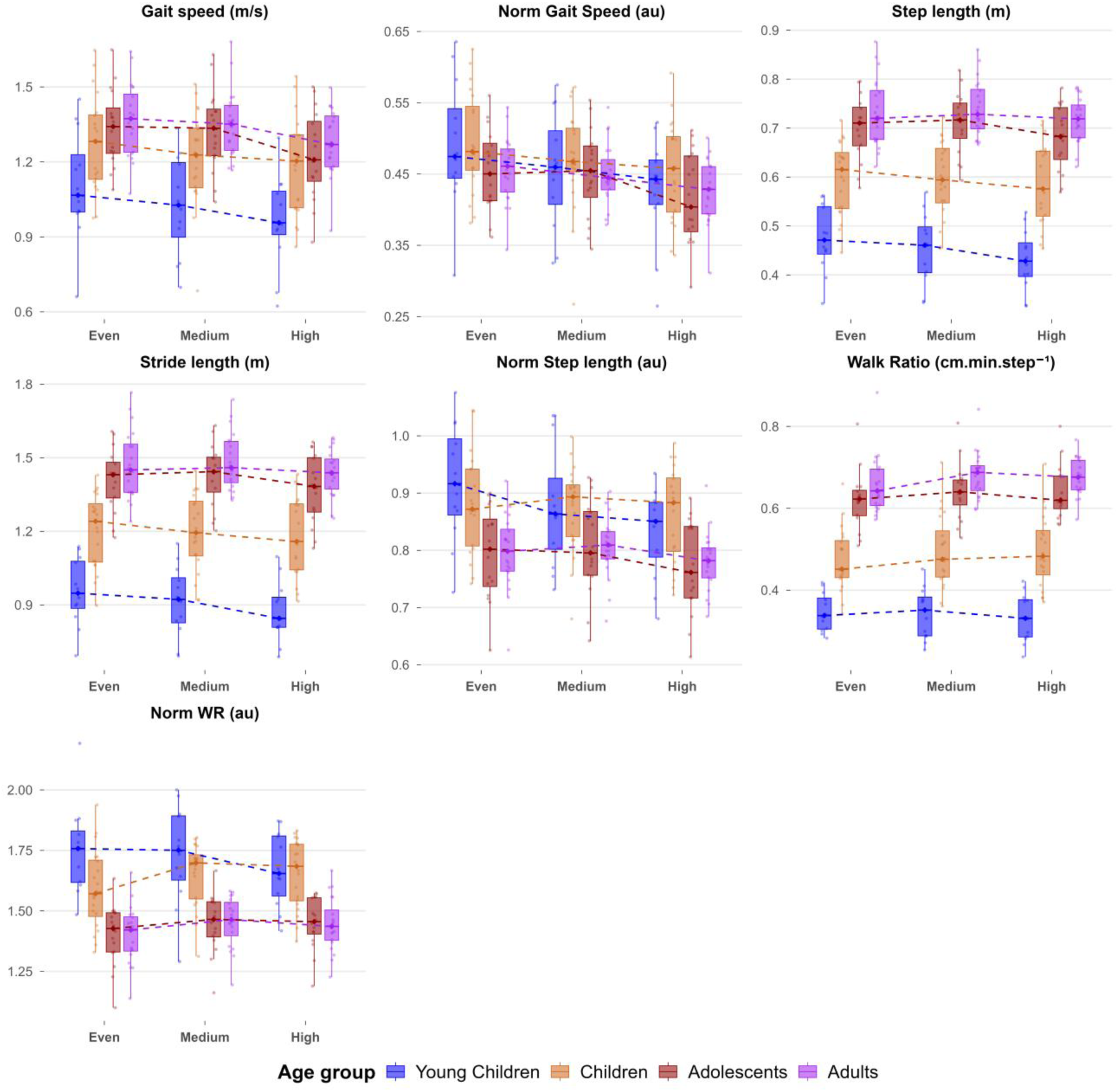
Pace-related variables presented for each age group (Young children, Children, Adolescents, and Adults) and surface (Even, Medium, and High surfaces). Boxplots represent the median (bold line), interquartile range (box limits), and 1.5 times the interquartile range (whiskers). Points represent individual participants’ data. *Abbreviations:* arbitrary unit (au); centimeter (cm); meter (m); minute (min); normalized (Norm); second (s); walk ratio (WR).

Significant main effect of Age was observed for all pace variables (q < 0.001, η²p [0.25-0.77]) except normalized gait speed (q = 0.32, η²p = 0.05). Gait speed, step and stride length, and walk ratio increased with Age, whereas normalized walk ratio decreased with Age. Post-hoc comparisons revealed significant differences between adults and both young children and children for all pace-related variables (p < 0.01, *d* [2.07-14.60]), except for gait speed, which significantly differed only between young children and adults (p < 0.001, *d* = 3.53). Finally, all pace variables decreased with surface irregularity increase (all q < 0.001, η²p [0.05-0.30]), except raw and normalized walk ratio that increased (q < 0.05, η²p = 0.12 and 0.05, respectively). Post-hoc comparison revealed significant differences between the even and high surfaces for step length and stride length (p < 0.001, *d* = 0.86 and 0.83, respectively), whereas the other variables showed significant differences between the even and medium surfaces (p < 0.05, *d* [0.45-0.70]).

#### Rhythm domain

Significant Age × Surface interaction was identified for SPARC (q < 0.001, η²p = 0.16). SPARC significantly decreased with increasing surface irregularity only in young children. On the even surface, young children showed significantly lower SPARC values than adolescents and adults (p < 0.05, *d* = 1.30 and 1.77, respectively). On uneven surfaces, this difference extended to all other age groups, with young children showing lower SPARC values than children, adolescents, and adults (p < 0.001, *d* = 2.84-3.77; Fig. 3).

**Figure 3:**
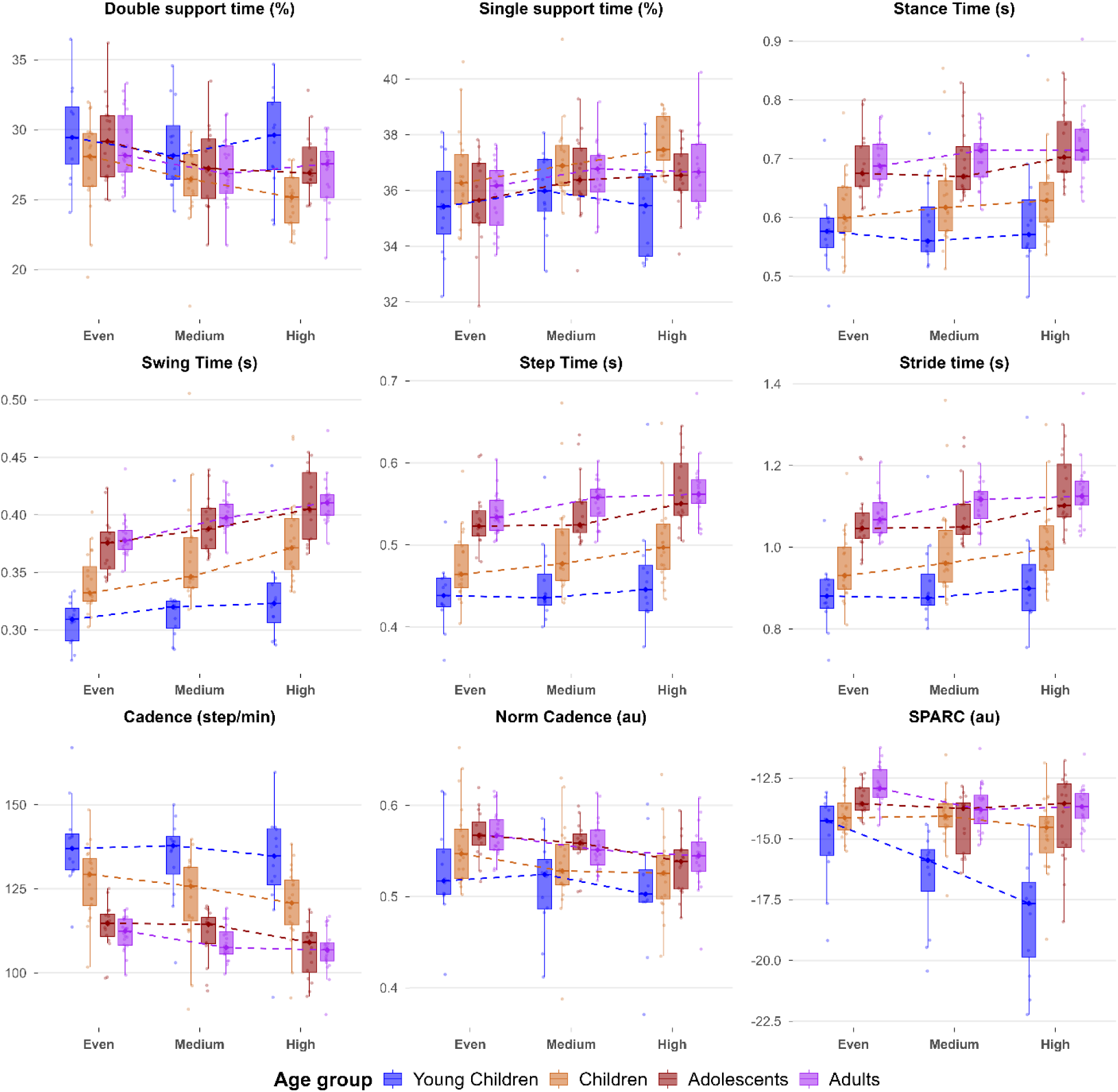
Rythm-related variables presented for each age group (Young children, Children, Adolescents, and Adults) and surface (Even, Medium, and High surfaces). Boxplots represent the median (bold line), interquartile range (box limits), and 1.5 times the interquartile range (whiskers). Points represent individual participants’ data. *Abbreviations:* arbitrary unit (au); minute (min); normalized (Norm); second (s); spectral arc length (SPARC).

Significant main effects of Age were observed for all rhythm variables (q < 0.001, η²p [0.14-0.55]). Cadence and double support time decreased with Age, whereas stance time, swing time, step time, stride time, and normalized cadence increased. Post-hoc comparisons indicated that rhythm-related variables differed significantly between adults and both young children and children (p < 0.001, *d* [1.87-5.01]), except for normalized cadence, for which young children differed significantly from adults (p = 0.02, *d* = 1.84). Then, all rhythm-related variables were significantly influenced by Surface irregularity (q < 0.001, η²p [0.14-51]). Double support time, raw and normalized cadence decreased whereas stance, swing, step, and stride time, and single support time increased with surface irregularity. Post-hoc comparison revealed that all these variables differed significantly between the even and uneven surfaces (medium and high surfaces) (p < 0.05, *d* [0.35-1.99]).

#### Dynamic stability domain

Significant Age × Surface interactions were observed for normalized step width (q = 0.02, η²p = 0.14), raw MoS AP (q = 0.01, η²p = 0.15), and normalized MoS AP (q < 0.001, η²p = 0.24). With increasing surface irregularity, normalized step width increased only in young children, who showed a significant difference between the even and high surfaces (p < 0.001, *d* = 1.55). On the even surface, young children had higher normalized step width than adolescents and adults (p < 0.01, *d* = 2.74 and 3.06, respectively). On the high surface, this difference extended to all other age groups, with young children showing higher values than children, adolescents, and adults (p < 0.001, *d* = 3.20-3.83) (Fig. 4). In contrast, raw and normalized MoS AP decreased in high surface only in young children and children groups (p < 0.001, *d* = 2.51 and 3.18 for raw and normalized variable respectively in young children, *d* = 1.07 and 1.02 for raw and normalized variable respectively in children). For raw MoS AP, this reduction led to a disappearance of the significant differences previously observed between young children and older groups on the even surface (p > 0.05), where young children exhibited higher values (Fig. 4). For normalized MoS AP, the decrease observed in young children and children resulted in significantly lower values in young children compared to all other age groups (p < 0.001, *d* [2.53-3.93]) and in children compared to adults (p < 0.01, *d* = 1.40) on high surface, whereas no such differences were present on the even surface.

**Figure 4:**
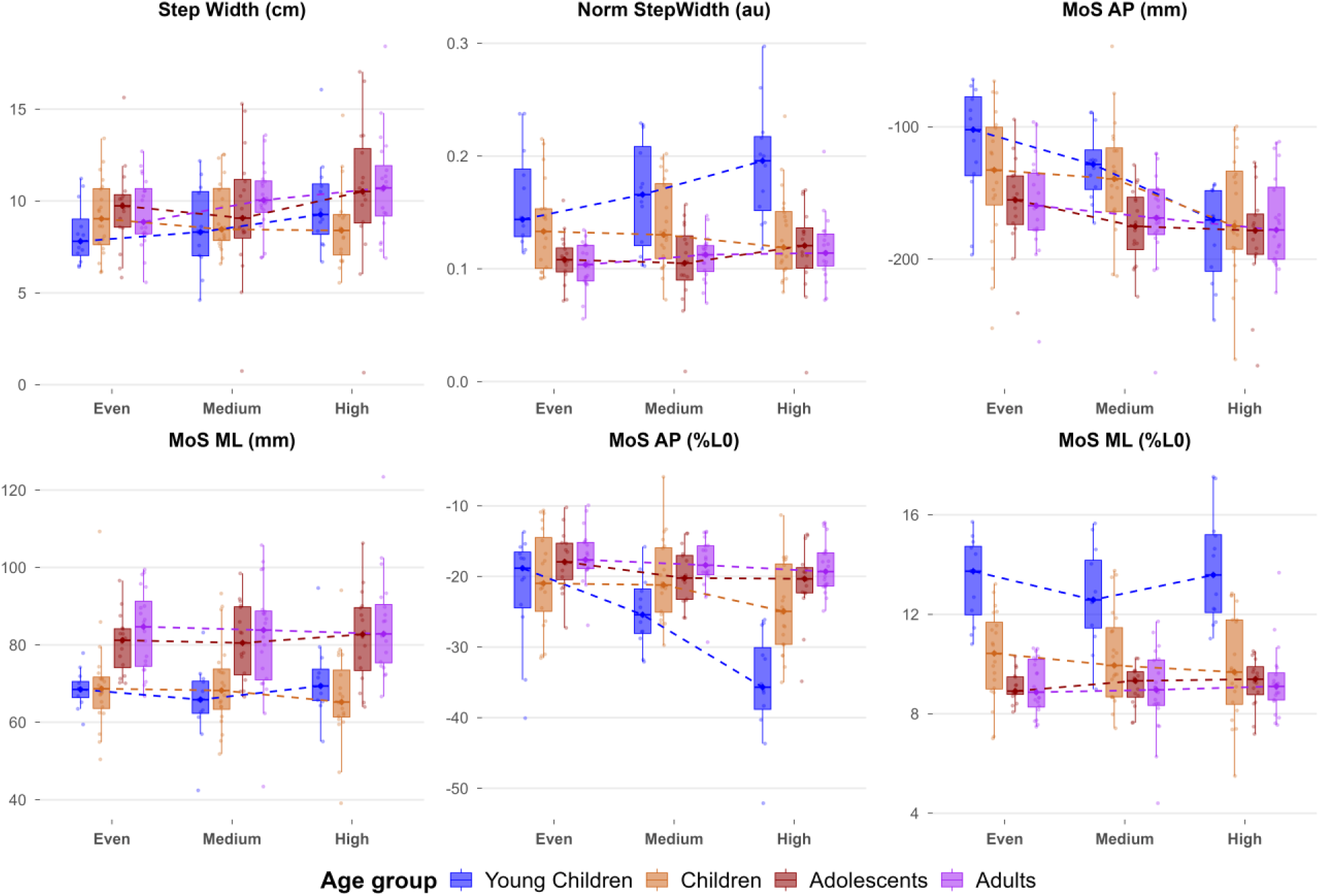
Dynamic-stability-related variables presented for each age group (Young children, Children, Adolescents, and Adults) and surface (Even, Medium, and High surfaces). Boxplots represent the median (bold line), interquartile range (box limits), and 1.5 times the interquartile range (whiskers). Points represent individual participants’ data. *Abbreviations:* anteroposterior (AP); arbitrary unit (au); centimeter (cm); leg length (L0); margin of stability (MoS); mediolateral (ML); millimeter (mm); normalized (Norm).

Significant main effects of Age were observed for raw and normalized MoS ML (q < 0.001, η²p = 0.34 and 0.54, respectively), but not for raw step width (q = 0.30, η²p = 0.05). Raw MoS ML increased with age, whereas normalized MoS ML decreased (Fig. 4). Post-hoc comparisons indicated that raw ML MoS differed significantly between adults and both young children and children (p < 0.001, *d* = 2.42 and 2.36, respectively). In contrast, normalized ML MoS differed significantly only between young children and adults (p < 0.001, *d* = 4.80). Then, significant main effects of Surface were observed for raw step width (q < 0.001, η²p = 0.09) with increasing values, but not for raw and normalized MoS ML (p = 0.26 and p = 0.15, respectively) (Fig. 4). Post-hoc comparisons showed that step width significantly differed only between the even and high surfaces (p < 0.001, *d* = 0.60).

#### Variability domain

Significant Age ×Surface interaction was identified for GVI (q < 0.001, η²p = 0.30). GVI decreased with increasing surface irregularity across all age groups, with a more pronounced reduction in adolescents and adults between even and high surfaces (p < 0.001, *d* = 3.78 and 3.88, respectively) compared to young children and children groups (p < 0.001, *d* = 1.23 and 1.53, respectively). On the even surface, all age groups differed significantly from one another (p < 0.01, *d* [1.20-4.58]), except between adolescents and adults who exhibited the highest values. With increasing surface irregularity these differences were attenuated, such that young children only differed from adolescents and adults on high surface (p < 0.01, *d* = 1.49 and 1.94, respectively), and children only from adults (p < 0.01, *d* = 1.04) (Fig. 5).

**Figure 5:**
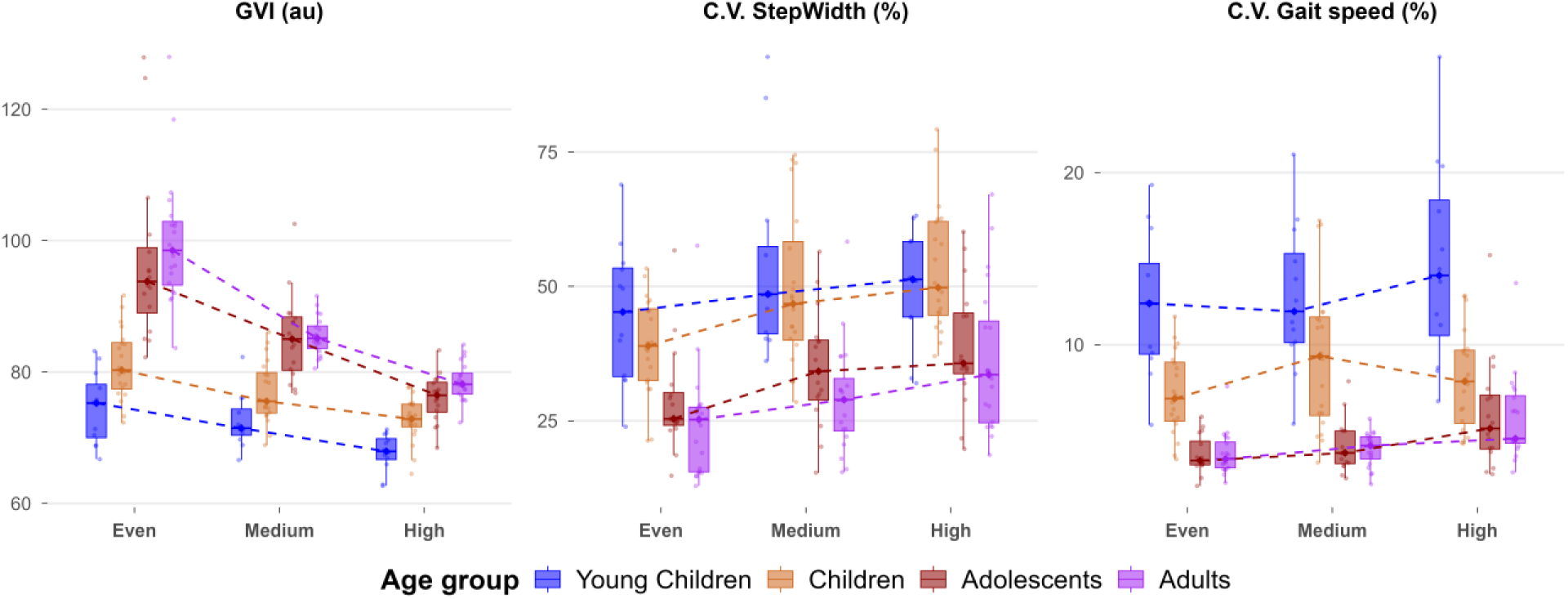
Variability-related variables presented for each age group (Young children, Children, Adolescents, and Adults) and surface (Even, Medium, and High). Boxplots represent the median (bold line), interquartile range (box limits), and 1.5 times the interquartile range (whiskers). Points represent individual participants’ data. *Abbreviations:* arbitrary unit (au); coefficient of variation (CV); gait variability index (GVI); normalized (Norm).

Significant main effects of Age were observed for CV step width and CV gait speed (q < 0.001, η²p = 0.43 and 0.65, respectively). Both decreased with Age (Fig. 5). Post-hoc comparisons indicated that both young children and children differed significantly from adults for CV step width (p < 0.001, *d* = 2.33 and 2.05, respectively) and CV gait speed (p < 0.001, *d* = 3.93 and 1.67, respectively). Then, significant main effects of Surface were reported for these two variables (q < 0.001, η²p = 0.29 and 0.16, respectively). Both increasing with surface irregularity (Fig. 5). Post-hoc comparisons revealed these variables significantly differed between even and uneven surfaces (medium and high surfaces) (p < 0.05, *d* [0.38-1.23]).

#### Asymmetry domain

Significant Age × Surface interactions were identified for SI stride length (q < 0.001, η²p = 0.22). SI stride length significantly increased with increasing surface irregularity only in young children, who were the only group showing significant difference between even and high surfaces (p < 0.001, *d* = 2.76) and significant differences with all other groups on the high surface (p < 0.001, *d* [2.75-2.87]) with higher values (Fig. 6).

**Figure 6:**
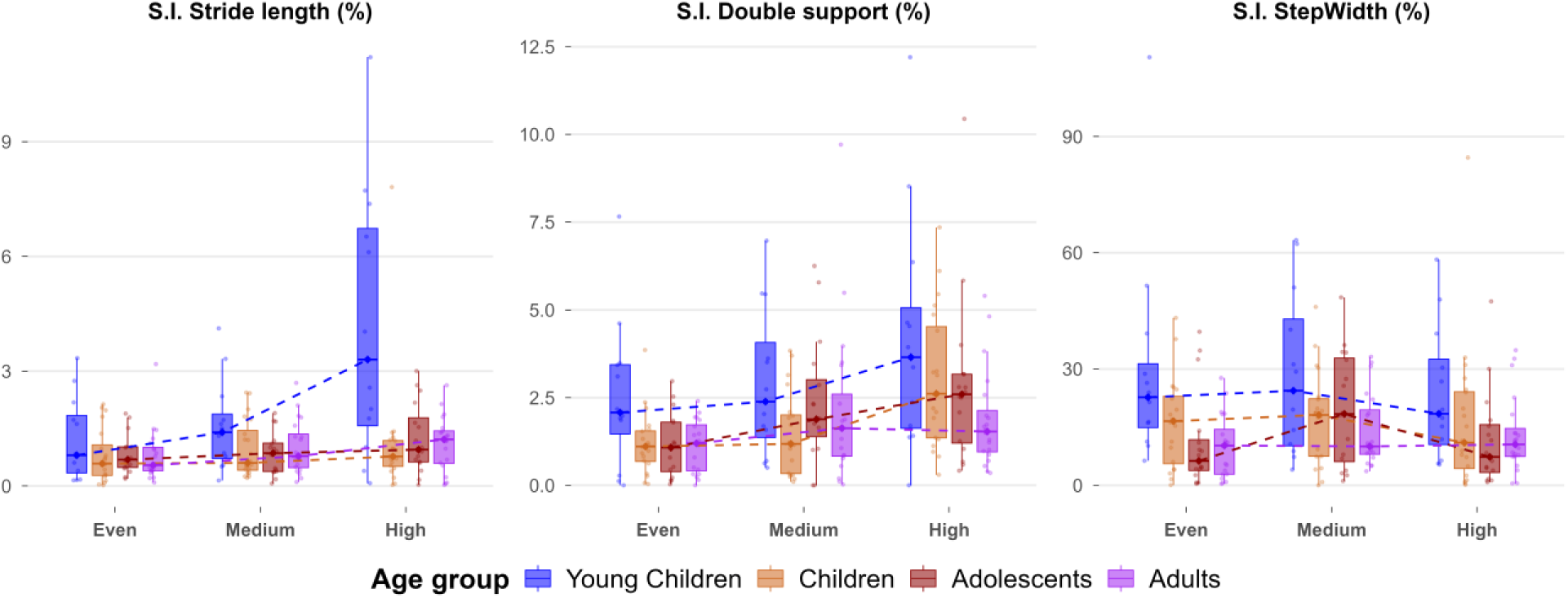
Asymmetry-related variables presented for each age group (Young children, Children, Adolescents, and Adults) and surface (Even, Medium, and High surfaces). Boxplots represent the median (bold line), interquartile range (box limits), and 1.5 times the interquartile range (whiskers). Points represent individual participants’ data. Abbreviations: normalized (Norm); symmetric index (SI).

Significant main effects of Age were observed for SI double support time and SI step width (q < 0.001, η²p = 0.08 and 0.28, respectively). Both decreased with age (Fig. 6). Post-hoc comparisons revealed that these variables differed significantly only between young children and adults (p < 0.001, *d* = 0.82 and 1.02, respectively). Then, significant main effects of Surface were observed for SI double support time (q < 0.001, η²p = 0.10), but not for SI step width (q = 0.33). SI double support time increased with surface irregularity (Fig. 6). Post-hoc comparisons revealed that SI double support time significantly differed between even and uneven surfaces (medium and high surfaces) (p ≤ 0.05, *d* = 0.39 and 0.81, for each surface respectively).

Age-specific timeline in the age at which gait variables differed from adults is summarized in Fig. 7.

**Figure 7:**
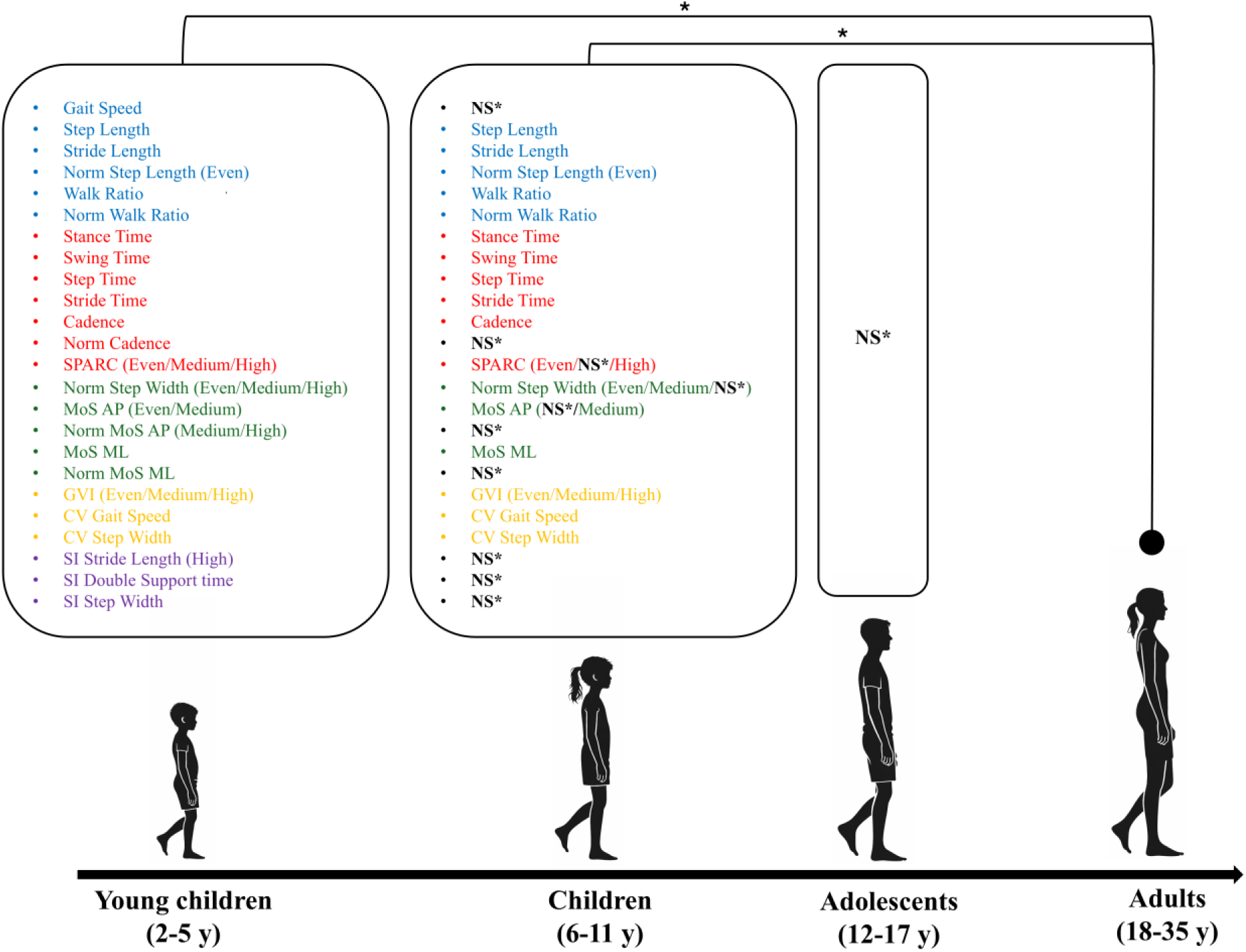
Developmental timeline illustrating the age at which gait variables differ from the Adult group (<0.05). Black labels and NS* indicate variables that were significantly different from adults in younger children but were no longer significantly different from adults in the corresponding age group. **Blue: Pace; Red: Rhythm; Green: Dynamic stability; Yellow: Variability; Purple: Asymmetry.** *Abbreviations:* antero-posterior (AP); arbitrary unit (au); centimeter (cm); coefficient of variation (CV); gait variability index (GVI); leg length (L0); meter (m); minute (min); medio-lateral (ML); millimeter (mm); margin of stability (MoS); normalized (Norm); second (s); symmetric index (SI); spectral arc length (SPARC).* Significant difference (p < 0.05) vs. adults.

### 3.3. Data-driven analysis

Four components were required to explain 70% of cumulative variance (Fig. 8a). PC1, PC2, PC3, and PC4 accounted for 32.4%, 20.0%, 10.1%, and 8.9% of the total variance, respectively, summing up 71.4% of explained variance across the 15 gait variables. The associations between PC are shown in supporting information (Fig. S1). Variables with absolute loadings superior to 0.50 were considered relevant (31). **PC1** was primarily driven by CV gait speed (0.84), normalized MoS ML (0.75), SPARC (-0.75), GVI (-0.75), normalized walk ratio (0.73), normalized step width (0.68), CV step width (0.57), normalized cadence (-0.55), and normalized MoS AP (-0.53). **PC2** was primarily driven by normalized gait speed (0.96), normalized step length (0.85), double support time (-0.69), normalized cadence (0.67), and normalized MoS AP (-0.52). **PC3** was primarily driven by SI double support time (0.52) and SI stride length (0.51). **PC4** was primarily driven by CV step width (0.64) and normalized step width (-0.51) (Fig. 8b). Using *k*-means clustering on the PCA-converted dataset, the optimal partition was obtained with two clusters, as supported by the within-cluster sum of squares (largest marginal reduction from k = 1 to k = 2: ΔWCSS = 695.92, 31.98%), the mean silhouette coefficient (k = 2: 0.51, higher than k = 3:10), and the adjusted rand index (k = 2: 0.85, higher than k = 3:10) (Fig. 8c). To visualize the retained two-cluster solution, observations were projected onto the first two principal components (Fig. 8d). Clusters characteristics are further detailed in the subsequent paragraphs.

**Figure 8:**
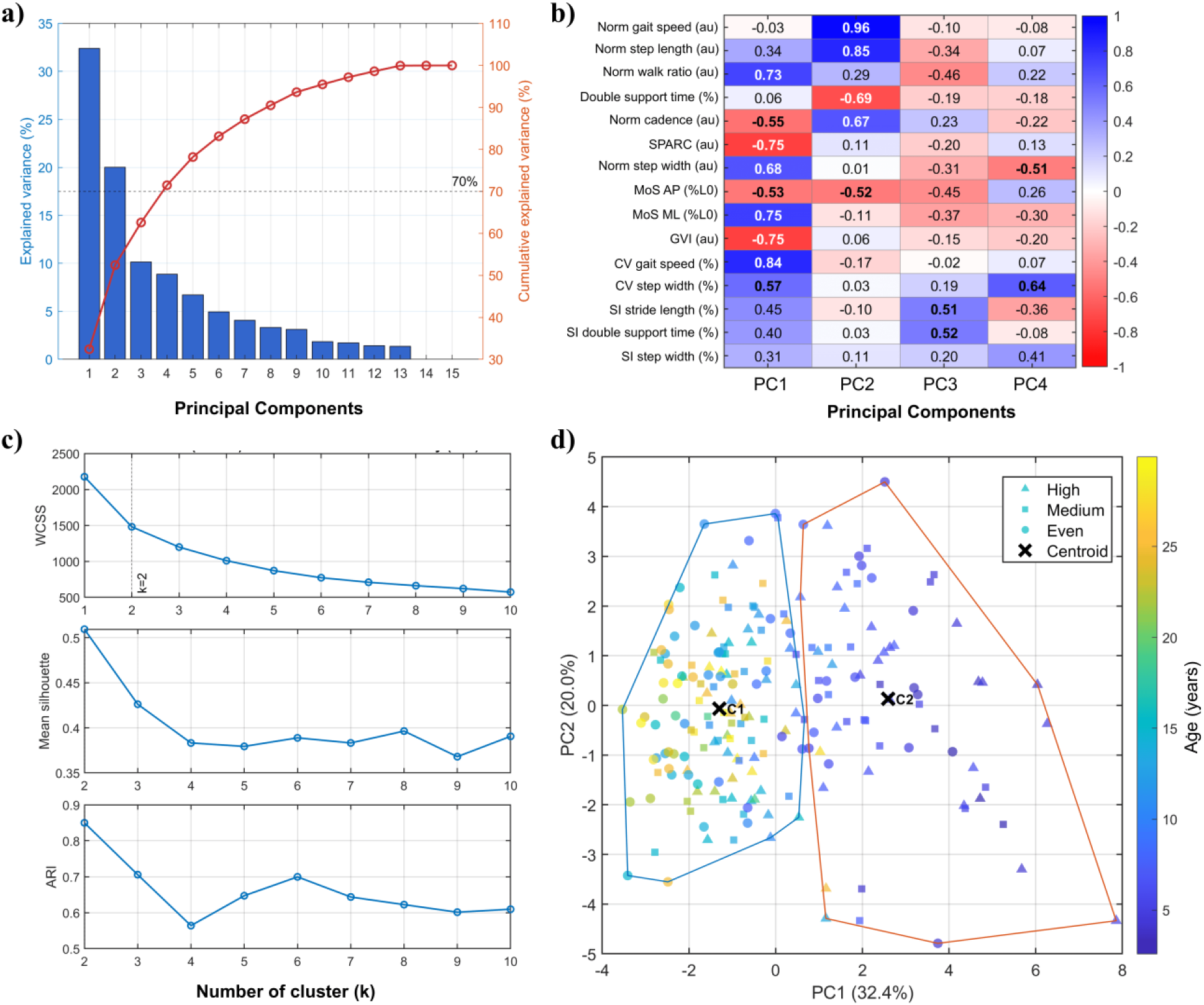
Principal component analysis and clustering results. **(a)** Scree plot showing the percentage of variance explained by each principal component (bars) and the cumulative explained variance (red curve). **(b)** Variable loadings for the four selected principal components. Color coding represents the magnitude and direction of each loading. **(c)** Cluster validation metrics computed for k = 1-10 clusters. The within-cluster sum of squares (WCSS) were assessed for k = 1-10, and partition quality and stability were evaluated for k ≥ 2 only. **(d)** PC1-PC2 score plot derived from z-standardized gait variables. Each point represents one participant-by-surface observation projected onto the first two principal component score dimensions. Points are color-coded according to age (y) and shape-coded according to surface irregularity. Convex hulls delineate cluster boundaries for k = 2. *Abbreviations:* antero-posterior (AP); adjusted rand index (ARI); arbitrary unit (au); coefficient of variation (CV); gait variability index (GVI); leg length (L0); medio-lateral (ML); margin of stability (MoS); normalized (Norm); principal component (PC); symmetric index (SI); spectral arc length (SPARC).

Cluster1 had more observations than Cluster2 (n=136 *vs.* n=68). Cluster1 comprised older participants (median and interquartile range: 182 [144-282] months) than Cluster2 (73 [55-87] months). Most adults and adolescents were assigned to Cluster1, representing 96.7% of adults (58/60) and 95.8% of adolescents (46/48). Conversely, most young children were assigned to Cluster2, representing 91.7% of this age group (33/36). Children were distributed more evenly between clusters, with 48.3% assigned to Cluster1 (29/60) and 51.7% to Cluster2 (31/60) (Fig. 9). Cluster distribution did not appear to vary according to sex (females: 67.4% vs. 65.3% in Clusters 1 and 2), but varied slightly across surfaces, with Cluster1 decreasing from Even to High (75.0% to 57.4%) and Cluster2 increasing accordingly (25.0% to 42.6%).

**Figure 9:**
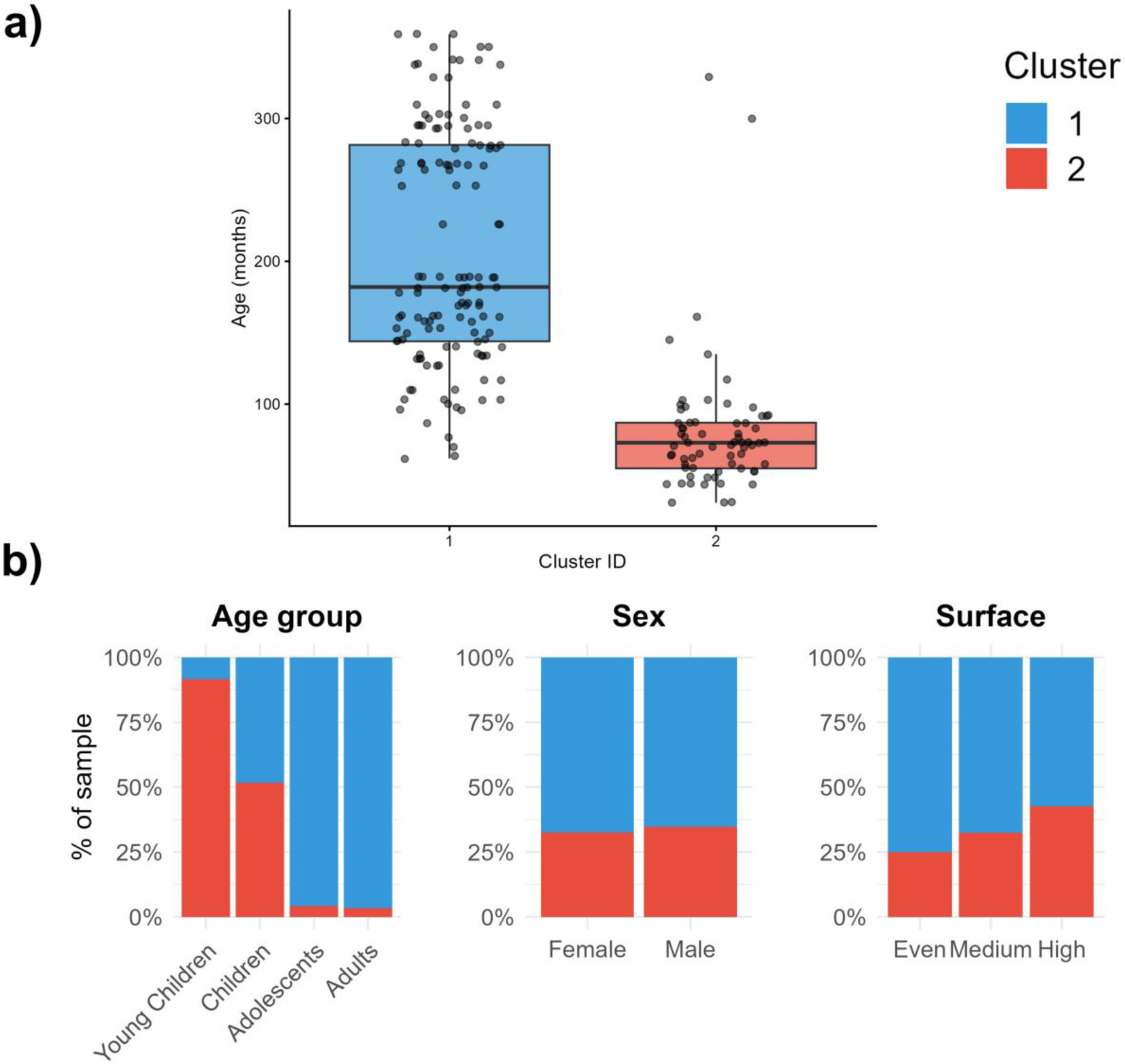
Cluster characteristics. **(a)** Distribution of age (in months) across the two clusters identified by *k*-means clustering. Boxplots represent the median (bold line), interquartile range (box limits), and 1.5 times the interquartile range (whiskers). Points represent individual participants’ data. **(b)** Proportion of participants assigned to each cluster according to age group (Young Children, Children, Adolescents, Adults), sex (Female, Male), and walking surface condition (Even, Medium, High). Blue: Cluster1; Red: Cluster2.

Across gait domains, pace and asymmetry was qualitatively higher in Cluster2 than Cluster1. Rhythm, tended to be smaller for Cluster2 than Cluster1. Finally, dynamic stability and variability depended of the variables. Cluster1 qualitatively showed lower normalized step width, MoS ML, and CV for gait speed and step width, together with higher GVI values and MoS AP. Normalized gait speed and double support time appeared similar between clusters (Fig. 10).

**Figure 10:**
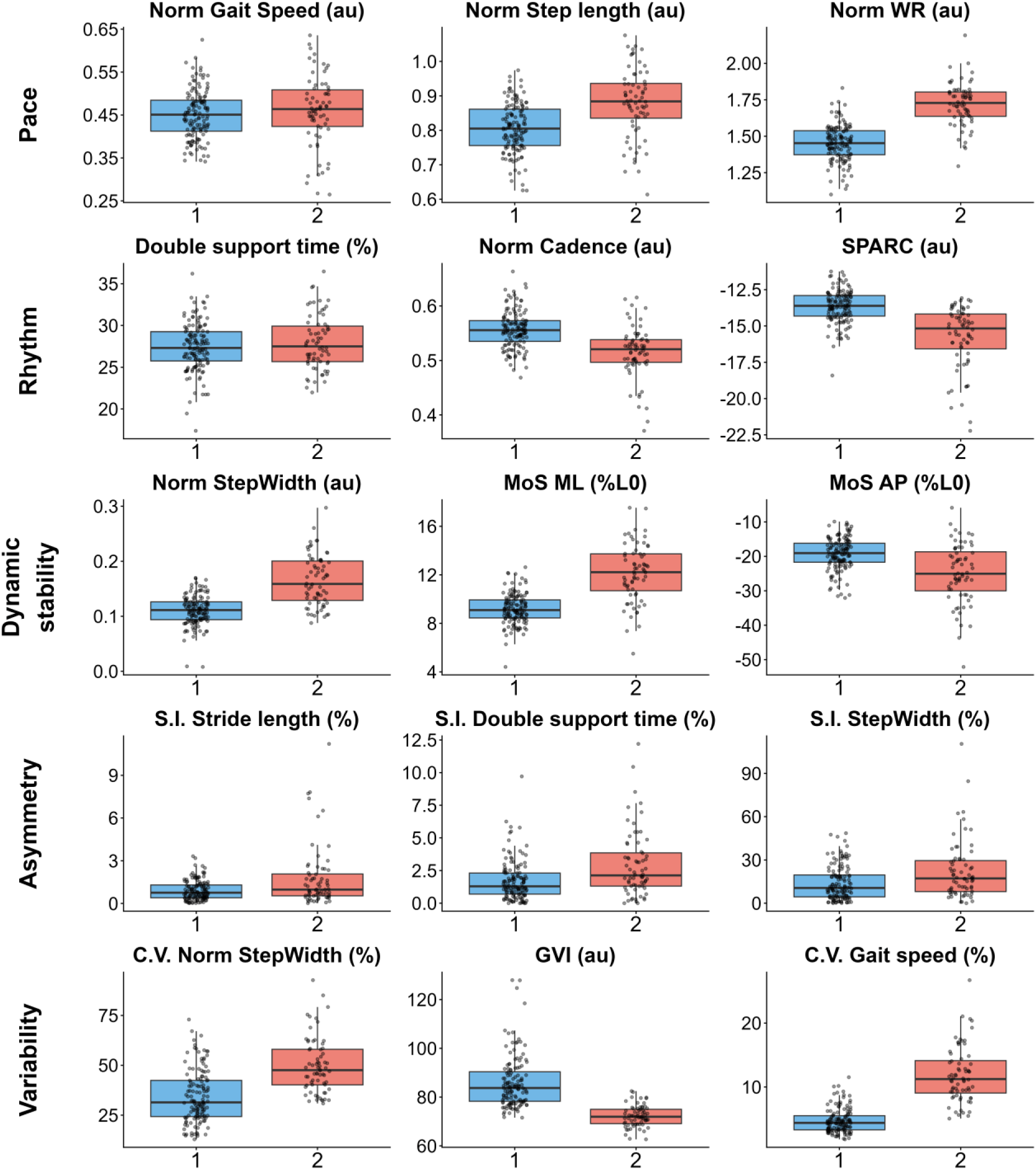
Gait variable distributions across clusters. Boxplots represent the median (bold line), interquartile range (box limits), and 1.5 times the interquartile range (whiskers). Points represent individual participants’ data. Blue: Cluster1; Red: Cluster2. *Abbreviations:* anteroposterior (AP); arbitrary unit (au); coefficient of variation (CV); gait variability index (GVI); leg length (L0); mediolateral (ML); margin of stability (MoS); normalized (Norm); symmetry index (SI); spectral arc length (SPARC); walk ratio (WR).

Further individual-level analysis revealed differences in cluster participants’ allocation across surfaces. Five subgroups were identified:

(i) *Consistent Cluster1*, participants who were consistently allocated to the Cluster1 across all surfaces (n = 38/68, 55.9%; 15.8 [13.2-23.6] years; 12M/26F);
(ii) *Consistent Cluster2,* participants who were consistently allocated to the Cluster2 across all surfaces (n = 15/68, 22.1%; 5.4 [4.1-6.6] years; 5M/10F).
(iii) *Early switchers,* participants who were allocated to the *Cluster1* on the even surface, but switched to the *Cluster2* on both uneven surfaces (n = 6/68, 8.8%, 7.7 [6.6-8.3] years; 3M/3F),
(iv) *Late switchers,* participants who were allocated to the Cluster1 on the even surface, but switched to the Cluster2 on High surface (n = 6/68, 8.8%, 12.8 [11.5-22.1] years; 2M/4F);
(v) *Transient,* participants, who exhibited inconsistent cluster allocation across surfaces (e.g., Cluster1 in Even, Cluster2 in Medium, then Cluster1 in High, or the reverse pattern) (n = 3/68, 4.4%; 5.8 [5.2-9.8] years; 2M/1F) (Fig.11);

**Figure 11:**
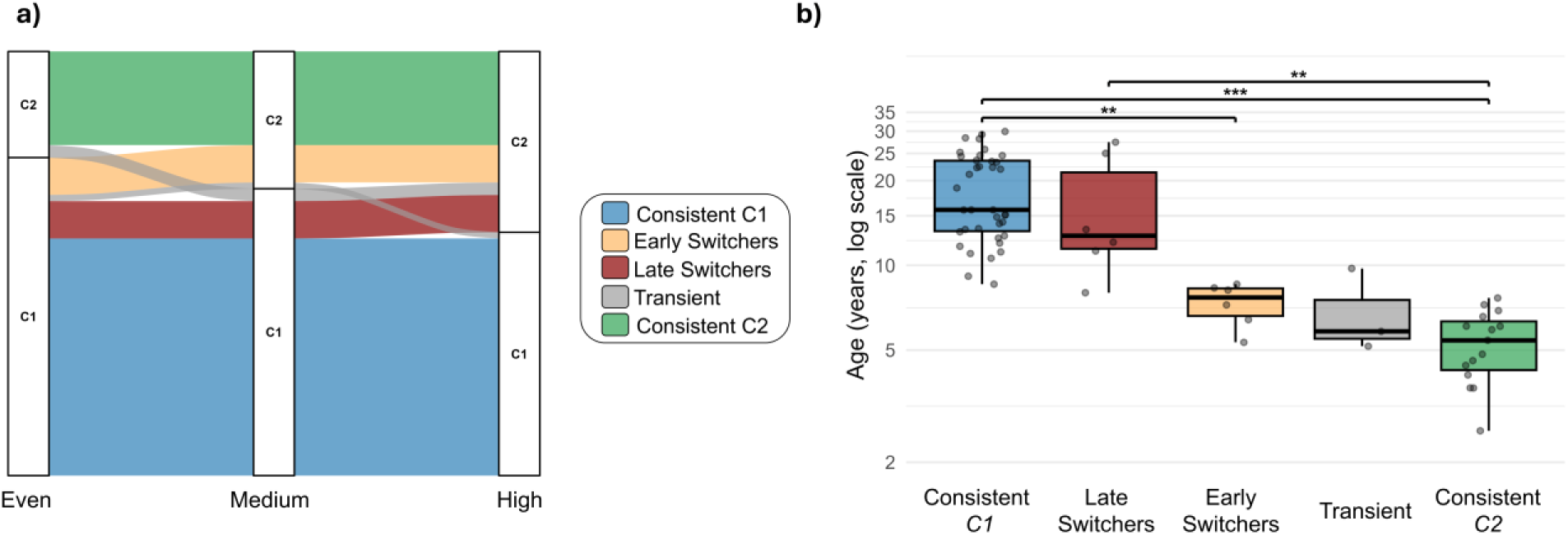
Profiles transitions across walking surface conditions. **(a)** Global alluvial representation of participants’ cluster allocation across Even, Medium, and High surface conditions. Subgroups are defined as follows: blue, *consistent in Cluster1* across all surfaces; light orange, *Early switchers* (participants transitioning from Cluster1 to the Cluster2 on the Medium surface); dark red, *Late switchers* (participants transitioning from Cluster1 to the Cluster2 on the High surface); grey, *Transient* (participants exhibiting non-systematic cluster transitions across surfaces); green, *consistent in Cluster2* across all surfaces. The width of each block reflects the proportion of participants within each cluster at each surface. **(b)** Boxplots represent the median (bold line), interquartile range (box limits), and 1.5 times the interquartile range (whiskers) of age (years) distribution across each subgroup. Points represent individual participants’ data and. Age is shown in years on a logarithmic y-axis for visualization only; statistics were performed on untransformed age values in months. ** indicates p < 0.01; *** indicates p < 0.001.

To explore age differences between these subgroups the Kruskal-Wallis test, followed by Dunn post-hoc comparisons with Holm correction, was used. Age differed significantly across these five subgroups (Kruskal-Wallis test, p < 0.001). Dunn’s post hoc comparisons with Holm correction revealed that the *consistent Cluster1* subgroup differed significantly from the *early switchers,* and *consistent Cluster2* subgroups (p < 0.01 and <0.001, respectively) and that *late switchers* differed significantly from the *consistent Cluster2* subgroup (p < 0.01) (Fig. 11). Sex distribution across these subgroups revealed no significant differences (Fisher’s exact test, p = 0.722).

To characterize the locomotor profiles of each subgroup, gait variables were qualitatively described across the three surface irregularity conditions, organized by domain (Fig. 12).

**Figure 12:**
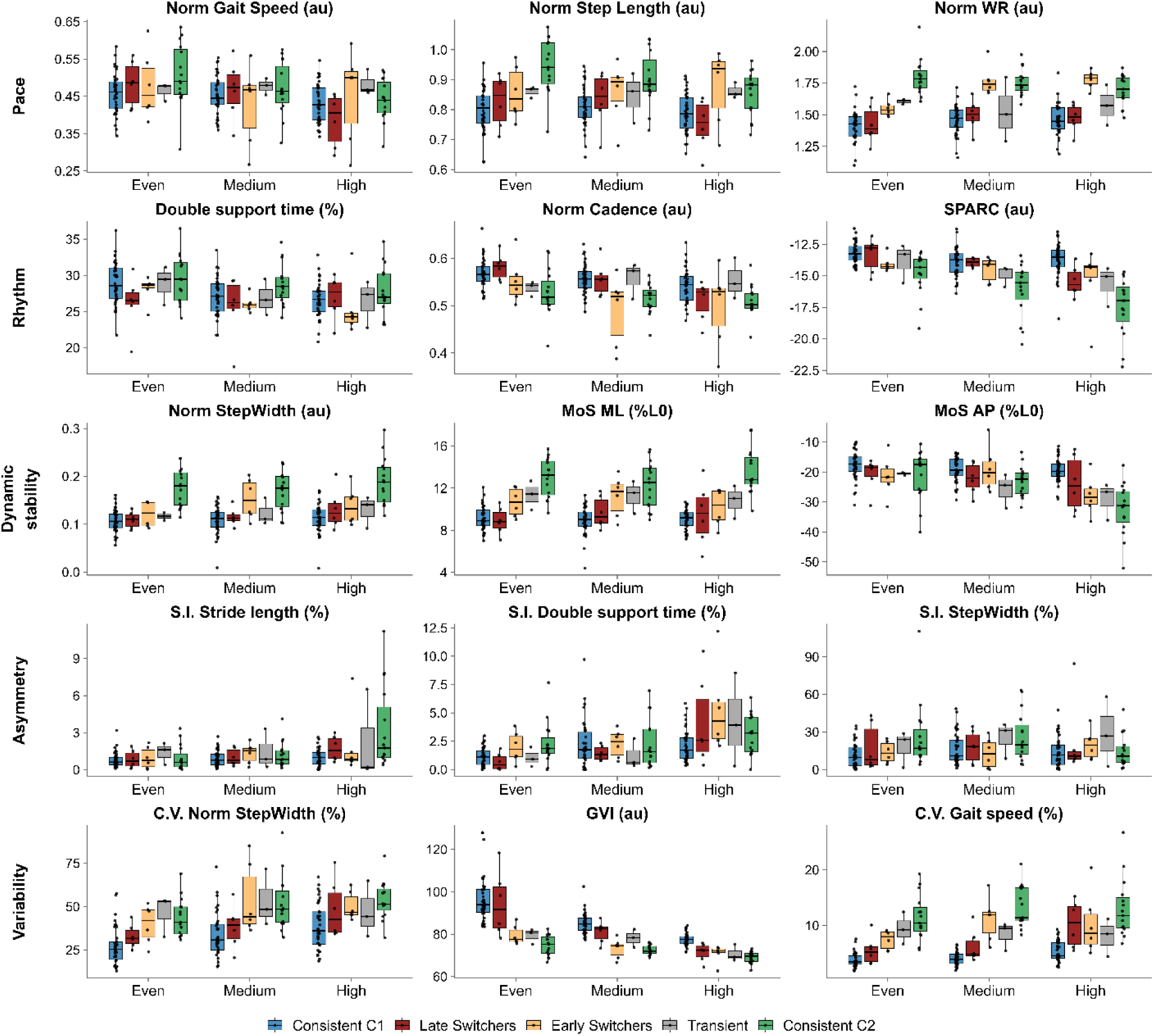
Gait variable distributions across subgroups and walking surface conditions. Boxplots represent the median (bold line), interquartile range (box limits), and 1.5 times the interquartile range (whiskers). Points represent individual participants’ data. *Abbreviations:* anteroposterior (AP); arbitrary unit (au); cluster1 (C1); cluster2 (C2); coefficient of variation (CV); gait variability index (GVI); leg length (L0); mediolateral (ML); margin of stability (MoS); normalized (Norm); symmetry index (SI); spectral arc length (SPARC); walk ratio (WR).

#### Pace domain

The *consistent Cluster2* subgroup exhibited the highest normalized step length and walk ratio across all conditions, whereas the *consistent Cluster1* subgroup consistently showed the lowest values. No clear differences emerged for normalized gait speed. *Early switchers*, *late switchers*, and *transient* presented intermediate values in an incremental order, with *late switchers* positioned closer to the *consistent Cluster1* subgroup. Surface irregularity did not produce a clear within-subgroup trend in pace variables. However, on uneven surfaces, *early switchers* diverged from the *consistent C1* subgroup and *late switchers*, displaying normalized step length and walk ratio values that approached those of the *consistent Cluster2* subgroup.

#### Rhythm domain

The *consistent Cluster1* subgroup exhibited the highest normalized cadence and SPARC values across conditions, whereas the *consistent Cluster2* subgroup generally showed the lowest values, particularly for SPARC on uneven surfaces. No clear pattern emerged for double support time. *Early Switchers, late switchers*, and *transient* presented intermediate values with the same incremental order described before. With increasing surface complexity, the *consistent Cluster2* subgroup exhibited a marked decrease in SPARC values relative to the other subgroups.

#### Dynamic stability domain

The *consistent Cluster2* subgroup exhibited higher normalized step width and normalized MoS ML values across all surfaces than the *consistent Cluster1* subgroup. Conversely, normalized MoS AP tended to be higher in the *consistent Cluster1* subgroup, particularly on the high surface, where lower values were observed in the *consistent Cluster2* subgroup and in *early switchers*. *Early Switchers, late switchers,* and *transient* presented intermediate values with the same incremental order described before for normalized MoS ML.

#### Asymmetry domain

The *consistent Cluster2* subgroup exhibited higher SI stride length on the high surface than the other subgroups, whereas the *consistent Cluster1* subgroup generally showed the lowest asymmetry values. SI double support time increased with surface irregularity across all subgroups except the *consistent Cluster1* subgroup. No clear surface-related pattern emerged for SI step width.

#### Variability domain

The *consistent Cluster1* subgroup exhibited the highest GVI values, followed by *late switchers*, whereas the *consistent Cluster2* subgroup and *early switchers* showed lower values. Conversely, CV of gait speed and CV of step width followed an opposite incremental pattern, increasing progressively from the *consistent Cluster1* subgroup toward the *consistent Cluster2* subgroup, with *late switchers*, *transient*, and *early switchers* presenting intermediate values in the same order described previously.

## 4. Discussion

This study aimed to characterize locomotor profiles across maturation using two complementary approaches: a hypothesis-driven approach evaluating the effects of age and surface complexity on multiple gait domains, and a data-driven clustering approach to identify locomotor profiles. Both approaches converged in showing that gait maturation is domain-specific and that locomotor profile expression is modulated by surface complexity, particularly in mid-childhood. These findings are discussed below in terms of their developmental significance and their mutual contribution to understanding locomotor maturation.

### Maturation of locomotion: learning from hypothesis-driven

The LMM analyses revealed a hierarchical progression of gait maturation that was not uniform across gait domains. Symmetry-related variables (SI double support time, SI step width) significantly differ only between the young children group (2-5 years) and adults, with large effect sizes (*d =* 0.82 and 1.02, respectively), suggesting an early stabilization of bilateral gait organization, consistent with previous studies (3,4). By contrast, gait variability (GVI, CV gait speed, CV step width), mediolateral dynamic stability (MoS ML), gait smoothness (SPARC), and pace (normalized walk ratio) related variables were significantly different between young children and adults as well as between children (6-11 years) adults (p < 0.05, *d >* 0.75), suggesting a more prolonged maturational trajectory for these gait domain. This prolonged trajectory, particularly for the mediolateral MoS, may reflect the greater control demands associated with mediolateral stability regulation compared with forward progression (34,35,52). While anteroposterior progression during walking partly relies on passive pendular dynamics, allowing the body to exploit gravity and passive mechanical exchanges, mediolateral stability depends more on active control of body motion and foot placement (52). Accordingly, the maturation of mediolateral MoS may depend on the progressive refinement of sensorimotor integration and anticipatory locomotor control. Importantly, the absence of a significant Age effect for normalized gait speed suggests that dimensionless forward progression is primarily governed by biomechanical scaling constraints rather than by progressive refinement of locomotor organization (32).

### Responses to surface complexity

Surface irregularity significantly affected nearly all gait variables across age groups (p < 0.05), confirming our first hypothesis. LMM analyses revealed that rhythm-related variables changed between already on the medium surface, whereas changes in pace and dynamic stability emerged only on the high surface, suggesting that the locomotor system initially regulates temporal flow to accommodate increased mechanical demands before adjusting step geometry. Despite this, mediolateral MoS was preserved across all surface conditions (η²p < 0.03), indicating active maintenance of lateral balance through step width adjustment rather than a degradation of frontal plane stability, consistent with the prioritization of mediolateral control as a primary determinant of fall prevention (35,53). This preservation was consistent across all age groups, suggesting that mediolateral dynamic stability represents a foundational constraint of bipedal locomotion rather than a maturational achievement, a competency that must be operational from the onset of independent walking (54). Age-specific perturbation provides further insight into developmental locomotion stages. Young children (2-5 years) exhibited the largest surface-induced changes across stride length asymmetry (η²p = 0.22), normalized step width (η²p = 0.14) and step length (η²p = 0.13), normalized MoS AP (η²p = 0.39), and gait smoothness (SPARC, η²p = 0.16), suggesting that surface irregularity imposes a greater functional challenge at this developmental stage. While direct comparisons are scarce in the literature, this pattern aligns with previous findings in children with cerebral palsy walking on irregular terrain, showing that uneven surfaces induce multidomain spatiotemporal adjustments across pace, rhythm, stability, and variability.

Some differences from typically developing peers were revealed only on uneven terrain, including greater stride width and increased gait speed variability (13). This supports the idea that irregular surfaces can expose locomotor-control constraints that remain less visible during even walking. In contrast, adolescents and adults demonstrated larger GVI reductions under increasing surface complexity (d = 2.08-3.88 *vs.* d = 0.86-1.53 in younger groups). This larger decrease may reflect a greater deviation from an initially more regular gait organization, whereas younger children already exhibited increased baseline variability on even surfaces, which may have limited the magnitude of further surface-induced modulation. This higher baseline variability may reflect a less stable locomotor organization during a developmental period in which perception-action coupling, multisensory integration, and anticipatory balance control are still being refined. Since irregular surfaces require visual detection of surface properties, anticipatory foot placement, and continuous recalibration of step geometry according to environmental affordances, these developmental constraints may have contributed to the larger surface-induced changes observed in young children (55–57). The present study extends this observation by showing that this higher baseline could limit further modulation when surface complexity increases.

### Refinement of locomotor profiles: learning from PCA-assisted clustering analysis

The switcher participants identified through clustering analysis refined the maturational timeline. The *k*-means algorithm identified two locomotor profiles. Based on the distinguishing gait features observed in the descriptive analysis and the main PC1 loadings, including CV gait speed (0.84), normalized MoS ML (0.75), SPARC (-0.75), and GVI (-0.75), Cluster1 (median age: 15.2 years) could be interpreted as a *smooth-regular profile*, characterized by higher gait smoothness, lower step-to-step variability, and a narrower base of support. Conversely, Cluster2 (median age: 6.1 years) could be interpreted as a *wide-base variable profile*, characterized by higher step-to-step variability, wider normalized step width, lower anteroposterior dynamic stability, and greater mediolateral stability margins. The variables driving cluster separation, primarily mediolateral stability, gait smoothness, and variability, correspond closely to the domains identified by the LMM as exhibiting the most protracted maturation, reinforcing their role as key dimensions of locomotor control organization.

The identification of 12 participants who transitioned from the *smooth-regular* to the *wide-base variable profile* as surface irregularity increased constitutes the most original contribution of the data-driven analysis (n = 12, 17.6%; median age: 8.5 [7.6-12.8] years). This surface-induced profile transition suggests that the Age × Surface interaction identified by LMM does not reflect isolated variable adjustments, but rather a coordinated reorganization of locomotor strategy involving simultaneous changes in variability, smoothness, base of support, and AP stability. The age gradient observed between *early* and *late switchers* (7.7 [6.6-8.3] and 12.8 [11.5-22.1] years, respectively) further suggests a progressive consolidation of locomotor robustness. Older switchers maintained the *smooth-regular profile* under moderate surface irregularity and shifted to the *wide-base variable profile* only under the most challenging condition. This interpretation is supported by the significant age differences observed across the five surface-transition subgroups. These findings provide a functional interpretation of the prolonged maturation of gait variability previously reported during even surface walking. Previous studies showed that GVI, and spatiotemporal variability continue to evolve beyond early childhood and stabilize only during late childhood or early adolescence (11-14 years) (9,58,59). Our findings extend these observations by showing that this developmental window is not only characterized by reduced variability during steady walking, but also by a progressive ability to preserve a *smooth-regular locomotor profile* when environmental constraints increase. This is consistent with split-belt treadmill findings showing that children up to 11 years exhibit slower rates of spatial locomotor adaptation than adults, supporting the later maturation of flexible locomotor control (59). Notably, children aged 6-11 years were distributed across both clusters (Cluster1: n = 29 observations; Cluster2: n = 31 observations), revealing substantial within-group heterogeneity that the predefined age-band structure of the LMM could not fully capture. Together, these findings suggest that the 7-13-year period is a transitional stage, during which gait becomes progressively more robust and less sensitive to surface complexity. Future studies should include larger samples within this transitional age range to determine whether locomotor maturation is better represented as a discrete two-profile structure or as a gradual continuum of intermediate profiles. Our results add a complementary and ecologically relevant perspective to previous evidence of prolonged gait maturation. Whereas earlier studies identified maturational differences under even or experimentally perturbed walking conditions (9,58,59), the present findings show that irregular surfaces can reveal context-dependent differences in locomotor organization during mid-childhood. Some children displayed a smooth-regular, adult-like profile on the even surface but shifted toward a wide-base variable profile when surface complexity increased, suggesting that locomotor maturity is not a fixed state but depends on task demands (60). From a clinical perspective, incorporating surface complexity may therefore complement standard even-surface gait assessments by revealing partially consolidated locomotor skills during this transitional developmental window.

The present study focused on global spatiotemporal and stability-related gait features, as these variables are widely used in clinical gait assessment and remain highly interpretable in relation to functional locomotor performance (61). However, this behavioral framework should be extended in future work by associating the maturation of underlying neural substrates and functional control strategies with locomotor profiles. The locomotor transitions identified here are particularly relevant within the 7-13 years window, as individuals in this age range showed context-dependent locomotor profiles. These gait profiles transitions may reflect the progressive development of an anticipatory locomotor control across childhood (62), which is fundamentally supported by the continuous maturation of multiple sensorimotor control processes. These processes include the progressive refinement of corticospinal pathways supporting distal motor control (63), the increasing complexity of neuromuscular modular organization (64), and the maturation of multisensory integration for balance (65,66). Future studies should combine complementary neurophysiological tools to determine which maturational processes contribute most directly to the consolidation of locomotor profiles. Anticipatory locomotor control could be assessed by combining kinematic and electromyography analyses to determine whether gaze behavior, foot placement and muscle activation are adjusted before or after contact with surface irregularities (55). Transcranial magnetic stimulation could assess corticospinal excitability and inhibition during gait-related phases, particularly in distal lower-limb muscles (67); electromyography could characterize muscle synergy organization, antagonist co-activation, and the temporal precision of muscle activation (64,68–70); and mobile high-density electroencephalography could quantify gait-related cortical dynamics (71,72). These measures could also be combined with spinal reflex assessments to characterize the maturation of spinal locomotor networks (73).

### Limitations and Perspectives

Several limitations should be acknowledged. First, because this study used a cross-sectional rather than a longitudinal design, it cannot capture within-individual developmental trajectories or directly track how locomotor responses evolve over time. Longitudinal studies will therefore be needed to better characterize the progression and stability of the locomotor profiles identified in the present study across development. Second, the number of participants within the transitional developmental range was limited (7-13 years, n = 19). This may have reduced the ability of the clustering procedure to directly detect intermediate profiles. Third, the uneven conditions differed from the even surface not only in geometric irregularity but also in material properties, as the Medium and High surfaces consisted of semi-rigid polyurethane panels while the even surface was rigid. Therefore, the observed gait responses may come from a combination of surface irregularity and surface smoothness. Nevertheless, irregular surfaced aimed at increasing the difficult to walk, which is exacerbated on semi-rigid surface. Finally, participants wore their own sport footwear, which may have introduced some uncontrolled variability related to shoe properties, although this also increased ecological validity by better reflecting everyday walking conditions.

## 5. Conclusion

Gait maturation appears to be primarily age-dependent, but its expression varies with task complexity, as younger children showing larger perturbations across smoothness, asymmetry, and stability gait domains on irregular surfaces. In contrast, adolescents and adults showed more selective adjustments, notably in gait variability, suggesting a more specific locomotor response to environmental constraints. The multivariate analysis identified two main locomotor profiles reflecting distinct levels of gait organization, with individuals in the ∼7-13 years range shifting between *smooth-regular* or *wide-base variable gait profiles* depending on surface complexity, indicating a critical period for the acquisition and consolidation of gait variability, smoothness, and dynamic stability control. Overall, these findings highlight the relevance of assessing gait under ecologically meaningful constraints and using multidimensional approaches to better characterize locomotor developmental profiles.

## Acknowledgments

The authors would like to thank Antoine Cohic, Sacha Douet, and Ines Toubal for their contribution to the investigation. We would like to thank the clinicians at Marie Enfant Rehabilitation Center for their assistance with recruitment.

## Author contributions

**SD:** Conceptualization; Data curation; Formal analysis; Investigation; Project administration; Visualization; Writing – Original draft preparation. **YC:** Conceptualization; Funding acquisition; Methodology; Project administration; Supervision; Validation; Writing – Review & Editing. **FDM:** Conceptualization; Supervision; Validation; Writing – Review & Editing. **ME:** Validation; Writing – Review & Editing.

## Data availability

Due to ethical restrictions at CHU Sainte-Justine, individual participant data cannot be shared publicly, as they may contain potentially identifying information. In addition, participants did not provide consent for their data to be deposited in a public repository. However, the statistical results are fully reported in the Supplementary Materials.

## Supporting information captions

**Table S1:** Descriptive values of variables per gait domain according to age group and walking surface.

**Table S2:** Effect sizes for all gait variables per domain derived from Linear Mixed Models.

**Table S3:** Post-hoc pairwise comparisons between age by surface for gait variables showing a significant effect of the interaction Age × Surface.

**Table S4:** Post-hoc pairwise comparisons between surface by age for gait variables showing a significant effect of the interaction Age × Surface.

**Table S5:** Post-hoc pairwise comparisons between each age group and adults for gait variables showing a significant effect of Age.

**Table S6:** Post-hoc pairwise comparisons between each irregular surface (Medium, High) and even for gait variables showing a significant effect of Surface.

**Figure S1:** Pairwise relationships between principal component (PC) scores derived from the global principal component analysis. Scatter plots display the distribution of observations across PC1-PC4, with color indicating age (years). Histograms along the diagonal represent the distribution of scores for each principal component. Percentages in parentheses denote the proportion of total variance explained by each component.

