## Supplementary files for "Maturation of Gait: Identification of Locomotor Profiles from Early Childhood to Adulthood"

### SUPPORTING INFORMATION

**Table S1:** Descriptive values of variables per gait domain according to age group and walking surface

| **Variable** | **Age groups** | | | | | | | | | | | | ***q-value**** | | |
|---|---|---|---|---|---|---|---|---|---|---|---|---|---|---|---|
| | **Young children** | | | **Children** | | | **Adolescents** | | | **Adults** | | | Age effect | Surface effect | Interaction |
| | *Even* | *Medium* | *High* | *Even* | *Medium* | *High* | *Even* | *Medium* | *High* | *Even* | *Medium* | *High* | | | |
| **Pace** | | | | | | | | | | | | | | | |
| **Gait speed (m/s)** | 1.11 [0.98-1.23] | 1.02 [0.91-1.14] | 0.96 [0.86-1.07] | 1.27 [1.19-1.36] | 1.21 [1.13-1.30] | 1.18 [1.09-1.26] | 1.34 [1.29-1.41] | 1.33 [1.25-1.41] | 1.22 [1.14-1.31] | 1.37 [1.30-1.43] | 1.35 [1.29-1.41] | 1.28 [1.22-1.34] | ***<0.001*** | ***<0.001*** | 0.69 |
| **Norm gait speed (au)** | 0.49 [0.44-0.54] | 0.45 [0.41-0.50] | 0.43 [0.38-0.47] | 0.49 [0.46-0.52] | 0.47 [0.44-0.50] | 0.46 [0.43-0.49] | 0.45 [0.43-0.48] | 0.45 [0.42-0.48] | 0.42 [0.38-0.45] | 0.46 [0.44-0.48] | 0.45 [0.43-0.47] | 0.43 [0.41-0.45] | 0.32 | ***<0.001*** | 0.45 |
| **Step length (m)** | 0.47 [0.44-0.51] | 0.45 [0.41-0.49] | 0.43 [0.39-0.47] | 0.60 [0.56-0.63] | 0.59 [0.56-0.63] | 0.59 [0.55-0.62] | 0.70 [0.67-0.73] | 0.71 [0.68-0.74] | 0.68 [0.65-0.71] | 0.73 [0.70-0.76] | 0.74 [0.71-0.76] | 0.72 [0.70-0.74] | ***<0.001*** | ***<0.001*** | 0.19 |
| **Norm step length (au)** | 0.92 [0.86-0.98] | 0.88 [0.82-0.94] | 0.83 [0.79-0.87] | 0.88 [0.84-0.92] | 0.87 [0.84-0.91] | 0.86 [0.83-0.90] | 0.79 [0.75-0.83] | 0.81 [0.76-0.85] | 0.77 [0.73-0.81] | 0.80 [0.77-0.83] | 0.81 [0.78-0.83] | 0.78 [0.76-0.80] | ***<0.001*** | ***<0.001*** | ***0.02*** |
| **Stride length (m)** | 0.95 [0.88-1.03] | 0.91 [0.83-1.00] | 0.87 [0.80-0.94] | 1.20 [1.14-1.27] | 1.19 [1.13-1.26] | 1.18 [1.11-1.25] | 1.41 [1.35-1.47] | 1.43 [1.37-1.49] | 1.37 [1.30-1.44] | 1.47 [1.41-1.53] | 1.49 [1.43-1.54] | 1.44 [1.40-1.48] | ***<0.001*** | ***<0.001*** | 0.29 |
| **Walk ratio** **($cm.min.pas^{-1}$)** | 0.35 [0.32-0.37] | 0.34 [0.31-0.38] | 0.33 [0.30-0.37] | 0.47 [0.44-0.51] | 0.49 [0.46-0.53] | 0.50 [0.46-0.54] | 0.62 [0.58-0.66] | 0.64 [0.61-0.68] | 0.64 [0.61-0.67] | 0.66 [0.63-0.69] | 0.68 [0.66-071] | 0.68 [0.66-0.70] | ***<0.001*** | ***<0.001*** | 0.14 |
| **Norm Walk ratio (au)** | 1.75 [1.65-1.86] | 1.73 [1.61-1.85] | 1.67 [1.58-1.76] | 1.59 [1.52-1.66] | 1.65 [1.59-1.71] | 1.65 [1.59-1.72] | 1.40 [1.34-1.46] | 1.45 [1.39-1.51] | 1.45 [1.40-1.50] | 1.40 [1.35-1.45] | 1.45 [1.41-1.50] | 1.44 [1.40-1.49] | ***<0.001*** | ***0.04*** | 0.08 |
| **Rythm** | | | | | | | | | | | | | | | |
| **Double support time (%)** | 29.73 [27.79-31.67] | 28.73 [26.91-30.54] | 29.12 [27.06-31.17] | 27.67 [26.27-29.06] | 26.35 [25.15-27.54] | 24.92 [24.08-25.75] | 29.12 [27.59-30.65] | 27.41 [26.00-28.83] | 27.41 [26.28-28.53] | 28.85 [27.74-29.96] | 27.07 [26.04-28.10] | 26.88 [25.79-27.97] | ***0.02*** | ***<0.001*** | 0.15 |
| **Single support time (%)** | 35.46 [34.46-36.47] | 35.99 [35.18-36.81] | 35.42 [34.39-36.46] | 36.50 [35.79-37.22] | 37.07 [36.48-37.66] | 37.74 [37.32-38.15] | 35.68 [34.88-36.48] | 36.48 [35.77-37.20] | 36.50 [35.91-37.09] | 35.82 [35.27-36.36] | 36.62 [36.13-37.11] | 36.80 [36.22-37.37] | ***0.02*** | ***<0.001*** | 0.09 |
| **Stance time (s)** | 0.58 [0.54-0.61] | 0.59 [0.55-0.63] | 0.60 [0.54-0.66] | 0.61 [0.58-0.64] | 0.63 [0.60-0.67] | 0.63 [0.60-0.66] | 0.69 [0.66-0.71] | 0.69 [0.66-0.72] | 0.72 [0.69-0.76] | 0.70 [0.68-0.71] | 0.70 [0.68-0.72] | 0.72 [0.70-0.75] | ***<0.001*** | ***<0.001*** | 0.79 |

#### Table S1 (continued)

| Variable | Age groups | | | | | | | | | | | | *q-value** | | |
|---|---|---|---|---|---|---|---|---|---|---|---|---|---|---|---|
| | **Young children** | | | **Children** | | | **Adolescents** | | | **Adults** | | | Age effect | Surface effect | Interaction |
| | *Even* | *Medium* | *High* | *Even* | *Medium* | *High* | *Even* | *Medium* | *High* | *Even* | *Medium* | *High* | | | |
| **Swing time (s)** | 0.31 [0.29-0.32] | 0.32 [0.30-0.34] | 0.33 [0.30-0.35] | 0.34 [0.33-0.35] | 0.37 [0.35-0.38] | 0.38 [0.36-0.39] | 0.37 [0.36-0.39] | 0.39 [0.38-0.40] | 0.41 [0.39-0.42] | 0.38 [0.37-0.39] | 0.40 [0.39-0.41] | 0.41 [0.40-0.42] | ***<0.001*** | ***<0.001*** | 0.75 |
| **Step time (s)** | 0.44 [0.42-0.46] | 0.45 [0.42-0.48] | 0.46 [0.42-0.50] | 0.48 [0.46-0.49] | 0.50 [0.47-0.53] | 0.50 [0.48-0.53] | 0.53 [0.51-0.55] | 0.54 [0.52-0.56] | 0.56 [0.54-0.59] | 0.54 [0.53-0.55] | 0.55 [0.54-0.56] | 0.57 [0.55-0.58] | ***<0.001*** | ***<0.001*** | 0.80 |
| **Stride time (s)** | 0.88 [0.83-0.93] | 0.91 [0.85-0.97] | 0.93 [0.85-1.01] | 0.95 [0.92-0.99] | 1.00 [0.95-1.05] | 1.01 [0.97-1.06] | 1.06 [1.03-1.10] | 1.08 [1.04-1.13] | 1.13 [1.09-1.18] | 1.08 [1.06-1.10] | 1.10 [1.08-1.13] | 1.13 [1.10-1.17] | ***<0.001*** | ***<0.001*** | 0.80 |
| **Cadence (step/min)** | 137.89 [130.35-145.43] | 133.98 [126.60-141.36] | 132.43 [122.99-141.86] | 127.15 [122.39-131.91] | 121.83 [116.13-127.53] | 120.15 [115.16-125.13] | 113.63 [110.02-117.24] | 111.29 [107.43-115.15] | 106.82 [102.79-110.86] | 111.61 [109.39-113.83] | 108.94 [106.61-111.27] | 106.47 [103.54-109.39] | ***<0.001*** | ***<0.001*** | 0.82 |
| **Norm cadence (au)** | 0.53 [0.50-0.56] | 0.51 [0.48-0.54] | 0.51 [0.47-0.54] | 0.56 [0.54-0.58] | 0.53 [0.51-0.55] | 0.53 [0.51-0.54] | 0.57 [0.55-0.58] | 0.56 [0.54-0.57] | 0.53 [0.52-0.55] | 0.57 [0.56-0.58] | 0.56 [0.54-0.57] | 0.54 [0.53-0.56] | ***0.02*** | ***<0.001*** | 0.80 |
| **SPARC (au)** | -14.93 [-15.99-(-13.87)] | -16.57 [-17.70-(-15.45)] | -18.18 [-19.54-(-16.82)] | -13.98 [-14.39-(-13.56)] | -14.13 [-14.63-(-13.62)] | -14.80 [-15.48-(-14.13)] | -13.38 [-13.71-(-13.06)] | -14.35 [-14.93-(-13.77)] | -14.18 [-15.10-(-13.27)] | -12.83 [-13.19-(-12.46)] | -13.72 [-14.15-(-13.29)] | -13.69 [-14.08-(-13.31)] | ***<0.001*** | ***<0.001*** | ***<0.001*** |
| | | | | | | | **Dynamic stability** | | | | | | | | |
| **Step width (cm)** | 8.23 [7.27-9.18] | 8.49 [7.17-9.81] | 9.85 [8.42-11.27] | 9.14 [8.24-10.05] | 9.20 [8.35-10.05] | 8.67 [7.71-9.62] | 9.64 [8.50-10.77] | 9.33 [7.53-11.12] | 10.51 [8.57-12.45] | 9.37 [8.54-10.19] | 10.12 [9.31-10.92] | 10.82 [9.60-12.04] | 0.30 | ***<0.001*** | 0.08 |
| **Norm step width (au)** | 0.16 [0.14-0.19] | 0.17 [0.14-0.19] | 0.19 [0.16-0.22] | 0.14 [0.12-0.15] | 0.14 [0.12-0.16] | 0.13 [0.11-0.15] | 0.11 [0.10-0.12] | 0.10 [0.09-0.12] | 0.12 [0.10-0.14] | 0.10 [0.09-0.11] | 0.11 [0.10-0.12] | 0.12 [0.10-0.13] | ***<0.001*** | ***<0.001*** | ***0.02*** |
| **MoS A.P. (mm)** | -113.96 [-139.61-(-88.30)] | -128.93 [-144.28-(-113.58)] | -181.49 [-201.52-(-161.46)] | -137.76 [-160.09-(-115.44)] | -138.55 [-157.11-(-119.98)] | -166.74 [-187.13-(-146.36)] | -157.15 [-174.04-(-140.27)] | -174.71 [-188.77-(-160.65)] | -183.59 [-203.25-(-163.93)] | -158.05 [-174.67-(-141.44)] | -169.03 [-185.43-(-152.63)] | -171.29 [-186.73-(-155.86)] | ***0.04*** | ***<0.001*** | ***0.01*** |
| **MoS M.L (mm)** | 68.56 [65.83-71.30] | 65.29 [59.73-70.86] | 70.97 [65.46-76.49] | 69.70 [64.29-75.11] | 69.73 [64.77-74.69] | 66.35 [61.18-71.53] | 80.36 [76.63-84.09] | 80.77 [75.92-85.63] | 82.09 [76.25-87.93] | 83.66 [79.03-88.29] | 81.61 [75.01-88.21] | 85.23 [79.50-90.96] | ***<0.001*** | 0.26 | 0.19 |

#### Table S1 (continued)

| Variable | Age groups | | | | | | | | | | | | *q-value** | | |
|---|---|---|---|---|---|---|---|---|---|---|---|---|---|---|---|
| | Young children | | | Children | | | Adolescents | | | Adults | | | Age effect | Surface effect | Interaction |
| | *Even* | *Medium* | *High* | *Even* | *Medium* | *High* | *Even* | *Medium* | *High* | *Even* | *Medium* | *High* | | | |
| **Norm MoS A.P (%L0)** | -21.81 [-26.42-(-17.20)] | -25.00 [-27.85-(-22.15)] | -35.39 [-39.74-(-31.04)] | -20.23 [-23.25-(-17.20)] | -20.34 [-22.87-(-17.81)] | -24.60 [-27.37-(-21.82)] | -17.88 [-19.95- (-15.80)] | -19.89 [-21.78-(-18.00)] | -20.92 [-23.51-(-18.33)] | -17.17 [-18.82-(-15.51)] | -18.38 [-20.06-(-16.69)] | -18.63 [-20.26-(-16.99)] | ***<0.001*** | ***<0.001*** | ***<0.001*** |
| **Norm MoS M.L (%L0)** | 13.39 [12.43-14.34] | 12.69 [11.55-13.83] | 13.83 [12.56-15.11] | 10.32 [9.52-11.12] | 10.34 [9.51-11.16] | 9.88 [8.94-10.81] | 9.08 [8.78-9.38] | 9.11 [8.72-9.51] | 9.26 [8.77-9.74] | 9.11 [8.62-9.59] | 8.91 [8.16-9.66] | 9.28 [8.66-9.90] | ***<0.001*** | 0.15 | 0.09 |
| **Variability** | | | | | | | | | | | | | | | |
| **GVI (au)** | 74.59 [71.51-77.66] | 72.53 [70.20-74.86] | 67.76 [66.16-69.36] | 81.22 [78.86-83.58] | 77.01 [74.94-79.08] | 72.76 [71.26-74.26] | 96.85 [90.43-103.27] | 85.31 [82.02-88.59] | 76.01 [74.21-77.81] | 100.00 [95.62-104.38] | 85.46 [84.19-86.74] | 78.51 [77.23-79.78] | ***<0.001*** | ***<0.001*** | ***<0.001*** |
| **CV gait speed (%)** | 12.26 [9.93-14.58] | 12.70 [10.26-15.13] | 14.66 [11.32-18.00] | 7.15 [6.11-8.19] | 9.36 [7.45-11.27] | 7.97 [6.70-9.24] | 3.62 [3.08-4.17] | 4.06 [3.32-4.81] | 5.93 [4.34-7.51] | 3.67 [3.13-4.21] | 3.93 [3.50-4.36] | 5.71 [4.62-6.80] | ***<0.001*** | ***<0.001*** | 0.08 |
| **CV step width (%)** | 44.77 [37.39-52.14] | 53.79 [43.63-63.96] | 50.11 [44.28-55.93] | 38.95 [34.97-42.92] | 49.46 [43.06-55.86] | 53.26 [48.07-58.45] | 28.72 [23.87-33.58] | 34.87 [29.48-40.26] | 38.77 [33.12-44.42] | 24.29 [19.72-28.87] | 29.16 [24.72-33.60] | 36.28 [30.26-42.30] | ***<0.001*** | ***<0.001*** | 0.29 |
| **Asymmetry** | | | | | | | | | | | | | | | |
| **SI stride length (%)** | 1.23 [0.61-1.84] | 1.56 [0.89-2.24] | 4.23 [2.25-6.22] | 0.79 [0.48-1.10] | 0.98 [0.65-1.31] | 1.12 [0.41-1.84] | 0.82 [0.57-1.08] | 0.84 [0.57-1.12] | 1.24 [0.79-1.69] | 0.81 [0.50-1.11] | 1.01 [0.70-1.31] | 1.11 [0.80-1.43] | ***<0.001*** | ***<0.001*** | ***<0.001*** |
| **SI double support time (%)** | 2.55 [1.33-3.77] | 2.91 [1.70-4.12] | 4.20 [2.24-6.16] | 1.19 [0.80-1.59] | 1.46 [0.94-1.98] | 3.01 [2.15-3.87] | 1.14 [0.69-1.59] | 2.36 [1.49-3.24] | 2.87 [1.67-4.08] | 1.12 [0.80-1.44] | 2.14 [1.14-3.14] | 1.87 [1.25-2.49] | ***<0.001*** | ***<0.001*** | 0.55 |
| **SI step width (%)** | 30.19 [14.16-46.22] | 28.50 [16.39-40.61] | 23.67 [13.87-33.48] | 16.12 [10.99-21.26] | 17.49 [12.20-22.77] | 16.72 [8.32-25.13] | 11.26 [5.00-17.51] | 19.58 [12.29-26.87] | 11.78 [5.59-17.97] | 9.92 [6.36-13.48] | 14.08 [9.92-18.23] | 13.08 [8.68-17.48] | ***<0.001*** | 0.33 | 0.80 |

***Notes:*** *Mean [95% Confidence Interval] are presented. *p-values were adjusted using the false discovery rate (FDR) correction (q-values).* ***Abbreviations:*** *anteroposterior (AP); arbitrary unit (au); centimeter (cm); coefficient of variation (CV); gait variability index (GVI); leg length (L0); meter (m); minute (min); mediolateral (ML); millimeter (mm); margin of stability (MoS); second (s); symmetry index (SI); spectral arc length (SPARC)*

**Commenté [FD1]:** Donc c'est plus des valeurs de p, c'est des valeurs de q.

**Table S2:** Effect sizes for all gait variables per domain derived from Linear Mixed Models.

| Variable | Effect | *F* | *p-value** | $\eta^2p$ | $R^2_m$ | $R^2_c$ |
|---|---|---|---|---|---|---|
| **Pace** | | | | | | |
| **Gait speed (m/s)** | Surface | 27.74 | ***<0.001*** | 0.30 | 0.30 | 0.83 |
| | Age | 9.82 | ***<0.001*** | 0.32 | | |
| | Surface*Age | 0.89 | 0.69 | 0.04 | | |
| **Norm gait speed (au)** | Surface | 27.33 | ***<0.001*** | 0.30 | 0.11 | 0.77 |
| | Age | 1.23 | 0.32 | 0.05 | | |
| | Surface*Age | 1.21 | 0.45 | 0.05 | | |
| **Step length (m)** | Surface | 14.96 | ***<0.001*** | 0.19 | 0.68 | 0.95 |
| | Age | 53.63 | ***<0.001*** | 0.72 | | |
| | Surface*Age | 1.79 | 0.19 | 0.08 | | |
| **Norm step length (au)** | Surface | 16.89 | ***<0.001*** | 0.21 | 0.24 | 0.83 |
| | Age | 6.93 | ***<0.001*** | 0.25 | | |
| | Surface*Age | 3.21 | **0.02** | 0.13 | | |
| **Stride length (m)** | Surface | 13.78 | ***<0.001*** | 0.18 | 0.68 | 0.95 |
| | Age | 53.10 | ***<0.001*** | 0.71 | | |
| | Surface*Age | 1.51 | 0.29 | 0.07 | | |
| **Walk ratio (cm.min.pas$^{-1}$)** | Surface | 8.75 | ***<0.001*** | 0.12 | 0.75 | 0.97 |
| | Age | 73.31 | ***<0.001*** | 0.77 | | |
| | Surface*Age | 2.07 | 0.14 | 0.09 | | |
| **Norm Walk ratio (au)** | Surface | 3.29 | **0.04** | 0.05 | 0.44 | 0.82 |
| | Age | 21.00 | ***<0.001*** | 0.50 | | |
| | Surface*Age | 2.53 | 0.08 | 0.11 | | |
| **Rythm** | | | | | | |
| **Double support time (%)** | Surface | 28.15 | ***<0.001*** | 0.31 | 0.19 | 0.78 |
| | Age | 3.45 | ***<0.001*** | 0.14 | | |
| | Surface*Age | 1.97 | 0.15 | 0.08 | | |
| **Single support time (%)** | Surface | 18.16 | ***<0.001*** | 0.22 | 0.18 | 0.74 |
| | Age | 3.77 | **0.02** | 0.15 | | |
| | Surface*Age | 2.33 | 0.09 | 0.10 | | |
| **Stance time (s)** | Surface | 10.20 | ***<0.001*** | 0.14 | 0.36 | 0.82 |
| | Age | 14.85 | ***<0.001*** | 0.41 | | |
| | Surface*Age | 0.67 | 0.79 | 0.03 | | |
| **Swing time (s)** | Surface | 65.40 | ***<0.001*** | 0.51 | 0.54 | 0.87 |
| | Age | 26.39 | ***<0.001*** | 0.55 | | |
| | Surface*Age | 0.77 | 0.75 | 0.03 | | |
| **Step time (s)** | Surface | 24.00 | ***<0.001*** | 0.27 | 0.45 | 0.85 |
| | Age | 20.79 | ***<0.001*** | 0.49 | | |
| | Surface*Age | 0.59 | 0.80 | 0.03 | | |

**Table S2 (Continued)**

| Variable | Effect | *F* | *p-value** | $\eta^2p$ | $R^2_m$ | $R^2_c$ |
|---|---|---|---|---|---|---|
| **Stride time (s)** | Surface | 24.54 | ***<0.001*** | 0.28 | 0.44 | 0.84 |
| | Age | 19.58 | ***<0.001*** | 0.48 | | |
| | Surface*Age | 0.57 | 0.80 | 0.03 | | |
| **Cadence (step/min)** | Surface | 22.45 | ***<0.001*** | 0.26 | 0.50 | 0.87 |
| | Age | 25.61 | ***<0.001*** | 0.55 | | |
| | Surface*Age | 0.48 | 0.82 | 0.02 | | |
| **Norm cadence (au)** | Surface | 26.04 | ***<0.001*** | 0.29 | 0.18 | 0.75 |
| | Age | 3.79 | **0.02** | 0.15 | | |
| | Surface*Age | 0.59 | 0.80 | 0.03 | | |
| **SPARC (au)** | Surface | 24.27 | ***<0.001*** | 0.27 | 0.46 | 0.59 |
| | Age | 28.42 | ***<0.001*** | 0.57 | | |
| | Surface*Age | 4.03 | ***<0.001*** | 0.16 | | |
| **Dynamic stability** | | | | | | |
| **Step width (cm)** | Surface | 6.41 | ***<0.001*** | 0.09 | 0.08 | 0.69 |
| | Age | 1.30 | 0.30 | 0.05 | | |
| | Surface*Age | 2.57 | 0.08 | 0.11 | | |
| **Norm step width (au)** | Surface | 6.86 | ***<0.001*** | 0.10 | 0.32 | 0.80 |
| | Age | 11.90 | ***<0.001*** | 0.36 | | |
| | Surface*Age | 3.34 | **0.02** | 0.14 | | |
| **MoS A.P. (mm)** | Surface | 27.17 | ***<0.001*** | 0.30 | 0.20 | 0.63 |
| | Age | 3.01 | **0.04** | 0.12 | | |
| | Surface*Age | 3.69 | **0.01** | 0.15 | | |
| **MoS M.L (mm)** | Surface | 1.40 | 0.26 | 0.02 | 0.29 | 0.78 |
| | Age | 11.05 | ***<0.001*** | 0.34 | | |
| | Surface*Age | 1.81 | 0.19 | 0.08 | | |
| **Norm MoS A.P (%L0)** | Surface | 29.86 | ***<0.001*** | 0.32 | 0.38 | 0.62 |
| | Age | 13.47 | ***<0.001*** | 0.39 | | |
| | Surface*Age | 6.63 | ***<0.001*** | 0.24 | | |
| **Norm MoS M.L (%L0)** | Surface | 2.01 | 0.15 | 0.03 | 0.48 | 0.85 |
| | Age | 25.31 | ***<0.001*** | 0.54 | | |
| | Surface*Age | 2.32 | 0.09 | 0.10 | | |
| **Variability** | | | | | | |
| **GVI (au)** | Surface | 110.46 | ***<0.001*** | 0.63 | 0.69 | 0.75 |
| | Age | 49.75 | ***<0.001*** | 0.70 | | |
| | Surface*Age | 9.13 | ***<0.001*** | 0.30 | | |
| **CV gait speed (%)** | Surface | 11.76 | ***<0.001*** | 0.16 | 0.55 | 0.76 |
| | Age | 40.31 | ***<0.001*** | 0.65 | | |
| | Surface*Age | 2.52 | 0.08 | 0.11 | | |
| **CV step width (%)** | Surface | 26.57 | ***<0.001*** | 0.29 | 0.39 | 0.70 |
| | Age | 16.18 | ***<0.001*** | 0.43 | | |
| | Surface*Age | 1.48 | 0.29 | 0.16 | | |

**Table S2 (Continued)**

| Variable | Effect | *F* | *p-value** | $\eta^2p$ | $R^2_m$ | $R^2_c$ |
|---|---|---|---|---|---|---|
| **Asymmetry** | | | | | | |
| **SI stride length (%)** | Surface | 16.06 | ***<0.001*** | 0.20 | *0.29* | *0.44* |
| | Age | 8.94 | ***<0.001*** | 0.30 | | |
| | Surface*Age | 6.19 | ***<0.001*** | 0.22 | | |
| **SI double support time (%)** | Surface | 10.71 | ***<0.001*** | 0.10 | *0.18* | *ø* |
| | Age | 5.64 | ***<0.001*** | 0.08 | | |
| | Surface*Age | 1.06 | 0.55 | 0.03 | | |
| **SI step width (%)** | Surface | 1.12 | 0.33 | 0.02 | *0.13* | *0.15* |
| | Age | 8.11 | ***<0.001*** | 0.28 | | |
| | Surface*Age | 0.54 | 0.80 | 0.02 | | |

***Notes:*** *$R^2m$ represents the variance explained by the fixed effects (Age, Surface, and their interaction), whereas $R^2c$ represents the variance explained by the full model, including the random effect (Participant). Estimates were derived from linear mixed-effects models fitted using restricted maximum likelihood (REML). Effect sizes for fixed factors were estimated using partial eta squared ($\eta^2p$), calculated using Satterthwaite's approximation for degrees of freedom. *p-values were adjusted using the false discovery rate (FDR) correction. Ø: For SI double support time, the conditional $R^2$ was not reported due to a singular fit, indicating negligible variance attributable to the random effect.* ***Abbreviations:*** *anteroposterior (AP); arbitrary unit (au); centimeter (cm); coefficient of variation (CV); gait variability index (GVI); leg length (L0); meter (m); minute (min); mediolateral (ML); millimeter (mm); margin of stability (MoS); second (s); symmetry index (SI); spectral arc length (SPARC).*

**Table S3:** Post-hoc pairwise comparisons between age by surface for gait variables showing a significant effect of the interaction Age × Surface.

| Variable | Comparison | Surface | Estimate [95%CI] | SE | df | t | *p-value** | *d* |
|---|---|---|---|---|---|---|---|---|
| **Pace** | | | | | | | | |
| **Normalized step length (au)** | YC vs. C | Even | 0.04 [-0.04-0.12] | 0.03 | 88.15 | 1.42 | 0.32 | *1.01* |
| | YC vs. Ado | | 0.12 [0.04-0.20] | 0.03 | 88.16 | 4.27 | **<0.001** | *3.28* |
| | YC vs. Adults | | 0.12 [0.04-0.20] | 0.03 | 88.00 | 4.16 | **<0.001** | *3.23* |
| | C vs. Ado | | 0.09 [0.01-0.16] | 0.03 | 88.16 | 3.31 | **<0.001** | *2.27* |
| | C vs. Adults | | 0.08 [0.02-0.15] | 0.03 | 87.80 | 3.16 | **<0.001** | *2.21* |
| | Ado vs. Adults | | -0.00 [-0.07-0.07] | 0.03 | 87.89 | -0.31 | 0.75 | *0.05* |
| | YC vs. C | Medium | 0.00 [-0.08-0.08] | 0.03 | 88.15 | 0.15 | 1 | *0.07* |
| | YC vs. Ado | | 0.07 [-0.01-0.15] | 0.03 | 88.16 | 2.43 | 0.07 | *1.88* |
| | YC vs. Adults | | 0.07 [-0.01-0.15] | 0.03 | 88.00 | 2.35 | 0.07 | *1.86* |
| | C vs. Ado | | 0.07 [-0.00-0.14] | 0.03 | 88.16 | 2.61 | 0.06 | *1.79* |
| | C vs. Adults | | 0.06 [0.00-0.14] | 0.02 | 87.80 | 2.53 | 0.07 | *1.81* |
| | Ado vs. Adults | | -0.00 [-0.07-0.07] | 0.02 | 87.89 | -0.20 | 1 | *0.02* |
| | YC vs. C | High | -0.03 [-0.11-0.05] | 0.03 | 88.15 | -1.09 | 0.56 | *0.84* |
| | YC vs. Ado | | 0.06 [-0.02-0.14] | 0.03 | 88.16 | 2.06 | 0.17 | *1.57* |
| | YC vs. Adults | | 0.04 [-0.03-0.13] | 0.03 | 88.00 | 1.61 | 0.33 | *1.34* |
| | C vs. Ado | | 0.09 [0.02-0.16] | 0.03 | 88.16 | 3.53 | **<0.01** | *2.42* |
| | C vs. Adults | | 0.08 [0.01-0.15] | 0.02 | 87.80 | 3.10 | **0.01** | *2.18* |
| | Ado vs. Adults | | -0.02 [-0.08-0.06] | 0.03 | 87.89 | -0.59 | 0.56 | *0.24* |

**Table S3 (Continued)**

| Variable | Comparison | Surface | Estimate [95%CI] | SE | df | t | *p-value** | *d* |
|---|---|---|---|---|---|---|---|---|
| **Rythm** | | | | | | | | |
| **SPARC (au)** | YC vs. C | Even | -0.94 [-2.28-0.38] | 0.50 | 170.93 | -1.90 | 0.18 | *0.80* |
| | YC vs. Ado | | -1.54 [-2.94-(-0.15)] | 0.52 | 170.95 | -2.96 | **0.02** | *1.30* |
| | YC vs. Adults | | -2.14 [-3.44-(-0.77)] | 0.50 | 170.59 | -4.30 | **<0.001** | *1.77* |
| | C vs. Ado | | -0.60 [-1.82-0.63] | 0.46 | 170.96 | -1.31 | 0.38 | *0.50* |
| | C vs. Adults | | -1.20 [-2.30-0.00] | 0.43 | 170.14 | -2.78 | **0.02** | *0.97* |
| | Ado vs. Adults | | -0.60 [-1.78-0.66] | 0.46 | 170.35 | -1.32 | 0.38 | *0.47* |
| | YC vs. C | Medium | -2.44 [-3.78-(-1.11)] | 0.50 | 170.93 | -4.90 | **<0.001** | *2.06* |
| | YC vs. Ado | | -2.22 [-3.62-(-0.83)] | 0.52 | 170.95 | -4.27 | **<0.001** | *1.87* |
| | YC vs. Adults | | -2.89 [-4.19-(-1.52)] | 0.50 | 170.59 | -5.81 | **<0.001** | *2.40* |
| | C vs. Ado | | 0.22 [-1.00-1.44] | 0.46 | 170.96 | 0.47 | 0.64 | *0.18* |
| | C vs. Adults | | -0.46 [-1.56-0.74] | 0.43 | 170.14 | -1.06 | 0.58 | *0.34* |
| | Ado vs. Adults | | -0.67 [-1.85-0.59] | 0.46 | 170.35 | -1.47 | 0.43 | *0.52* |
| | YC vs. C | High | -3.36 [-4.71-(-2.04)] | 0.50 | 170.93 | -6.76 | **<0.001** | *2.84* |
| | YC vs. Ado | | -3.99 [-5.39-(-2.60)] | 0.52 | 170.95 | -7.67 | **<0.001** | *3.36* |
| | YC vs. Adults | | -4.52 [-5.81-(-3.15)] | 0.50 | 170.59 | -9.08 | **<0.001** | *3.77* |
| | C vs. Ado | | -0.62 [-1.81-0.60] | 0.46 | 170.96 | -1.37 | 0.35 | *0.52* |
| | C vs. Adults | | -1.15 [-2.26-0.04] | 0.43 | 170.14 | -2.68 | **0.02** | *0.93* |
| | Ado vs. Adults | | -0.54 [-0.49-0.74] | 0.46 | 170.35 | -1.17 | 0.35 | *0.41* |

**Table S3 (Continued)**

| Variable | Comparison | Surface | Estimate [95%CI] | SE | df | t | *p-value** | *d* |
|---|---|---|---|---|---|---|---|---|
| **Dynamic stability** | | | | | | | | |
| **Normalized step width (au)** | YC vs. C | Even | 0.03 [-0.01-0.06] | 0.01 | 94.25 | 1.89 | 0.12 | *1.28* |
| | YC vs. Ado | | 0.05 [0.02-0.09] | 0.01 | 94.26 | 3.87 | **<0.01** | *2.74* |
| | YC vs. Adults | | 0.06 [0.02-0.10] | 0.01 | 94.05 | 4.53 | **<0.001** | *3.06* |
| | C vs. Ado | | 0.03 [-0.00-0.06] | 0.01 | 94.26 | 2.34 | 0.06 | *1.46* |
| | C vs. Adults | | 0.04 [0.00-0.07] | 0.01 | 93.81 | 3.04 | **0.01** | *1.77* |
| | Ado vs. Adults | | 0.01 [-0.03-0.04] | 0.01 | 93.93 | 0.54 | 0.59 | *0.31* |
| | YC vs. C | Medium | 0.03 [-0.01-0.06] | 0.01 | 94.25 | 2.06 | 0.09 | *1.39* |
| | YC vs. Ado | | 0.06 [0.02-0.10] | 0.01 | 94.26 | 4.43 | **<0.001** | *3.14* |
| | YC vs. Adults | | 0.06 [0.02-0.09] | 0.01 | 94.06 | 4.20 | **<0.001** | *2.84* |
| | C vs. Ado | | 0.03 [0.00-0.07] | 0.01 | 94.26 | 2.81 | **0.02** | *1.75* |
| | C vs. Adults | | 0.03 [-0.00-0.06] | 0.01 | 93.81 | 2.47 | **0.05** | *1.44* |
| | Ado vs. Adults | | -0.00 [-0.04-0.03] | 0.01 | 93.93 | -0.46 | 0.64 | *0.31* |
| | YC vs. C | High | 0.06 [0.03-0.10] | 0.01 | 94.25 | 4.72 | **<0.001** | *3.20* |
| | YC vs. Ado | | 0.07 [0.04-0.11] | 0.01 | 94.26 | 5.40 | **<0.001** | *3.83* |
| | YC vs. Adults | | 0.08 [0.04-0.11] | 0.01 | 94.06 | 5.40 | **<0.001** | *3.83* |
| | C vs. Ado | | 0.01 [-0.02-0.04] | 0.01 | 94.26 | 1.00 | 0.84 | *0.62* |
| | C vs. Adults | | 0.01 [-0.02-0.04] | 0.01 | 93.81 | 1.09 | 0.84 | *0.62* |
| | Ado vs. Adults | | 0.00 [-0.03-0.03] | 0.01 | 93.93 | 0.03 | 0.98 | *0.00* |

#### Table S3 (Continued)

| Variable | Comparison | Surface | Estimate [95%CI] | SE | df | t | *p-value** | *d* |
|---|---|---|---|---|---|---|---|---|
| **Dynamic stability** | | | | | | | | |
| **MoS AP (mm)** | YC vs. C | Even | 23.38 [-14.96-62.57] | 14.35 | 120.86 | 1.63 | 0.32 | *0.88* |
| | YC vs. Ado | | 42.93 [2.66-83.73] | 15.00 | 120.88 | 2.86 | **0.02** | *1.61* |
| | YC vs. Adults | | 45.60 [5.33-82.86] | 14.38 | 120.54 | 3.17 | **0.01** | *1.64* |
| | C vs. Ado | | 19.55 [-16.21-55.00] | 13.18 | 120.12 | 1.78 | 0.31 | *0.72* |
| | C vs. Adults | | 22.22 [-13.28-53.86] | 12.49 | 120.12 | 1.78 | 0.31 | *0.75* |
| | Ado vs. Adults | | 2.67 [-34.71-36.51] | 13.23 | 120.31 | 0.20 | 0.84 | *0.03* |
| | YC vs. C | Medium | 9.18 [-29.15-48.38] | 14.35 | 120.86 | 0.64 | 1 | *0.36* |
| | YC vs. Ado | | 45.50 [5.23-86.31] | 15.00 | 120.88 | 3.03 | **0.02** | *1.70* |
| | YC vs. Adults | | 41.60 [1.33-78.86] | 14.38 | 120.54 | 2.89 | **0.02** | *1.49* |
| | C vs. Ado | | 36.32 [0.55-71.76] | 13.18 | 120.89 | 2.76 | **0.03** | *1.35* |
| | C vs. Adults | | 32.42 [-3.09-64.05] | 12.49 | 120.12 | 2.60 | **0.03** | *1.13* |
| | Ado vs. Adults | | -3.90 [-41.28-29.93] | 13.23 | 120.31 | -0.29 | 1 | *0.21* |
| | YC vs. C | High | -15.18 [-53.51-24.01] | 14.35 | 120.86 | -1.06 | 1 | *0.55* |
| | YC vs. Ado | | 1.83 [-38.44-42.64] | 15.00 | 120.88 | 0.12 | 1 | *0.08* |
| | YC vs. Adults | | -8.69 [-48.96-28.57] | 14.38 | 120.54 | -0.60 | 1 | *0.38* |
| | C vs. Ado | | 17.00 [-18.76-52.45] | 13.18 | 120.89 | 1.29 | 1 | *0.63* |
| | C vs. Adults | | 6.49 [-29.02-38.12] | 12.49 | 120.12 | 0.52 | 1 | *0.17* |
| | Ado vs. Adults | | -10.52 [-47.90-23.31] | 13.23 | 120.31 | -0.80 | 1 | *0.46* |

**Table S3 (Continued)**

| Variable | Comparison | Surface | Estimate [95%CI] | SE | df | t | *p-value** | *d* |
|---|---|---|---|---|---|---|---|---|
| **Dynamic stability** | | | | | | | | |
| **Normalized MoS AP (%L0)** | YC vs. C | Even | -1.66 [-6.89-3.73] | 1.94 | 150.29 | -0.85 | 0.79 | *0.37* |
| | YC vs. Ado | | -3.98 [-9.49-1.62] | 2.03 | 150.31 | -1.95 | 0.26 | *0.92* |
| | YC vs. Adults | | -4.36 [-9.96-0.67] | 1.95 | 149.89 | -2.23 | 0.16 | *1.09* |
| | C vs. Ado | | -2.32 [-7.23-2.53] | 1.78 | 150.32 | -1.29 | 0.59 | *0.55* |
| | C vs. Adults | | -2.69 [-7.67-1.54] | 1.69 | 149.38 | -1.59 | 0.45 | *0.72* |
| | Ado vs. Adults | | -0.37 [-5.59-4.17] | 1.79 | 149.62 | -0.21 | 0.83 | *0.17* |
| | YC vs. C | Medium | -4.73 [-9.97-0.65] | 1.94 | 150.29 | -2.43 | 0.06 | *1.09* |
| | YC vs. Ado | | -5.15 [-10.67-0.45] | 2.03 | 150.31 | -2.53 | 0.06 | *1.20* |
| | YC vs. Adults | | -6.34 [-11.94-(-1.31)] | 1.95 | 149.89 | -3.24 | **<0.01** | *1.55* |
| | C vs. Ado | | -0.41 [-5.33-4.43] | 1.78 | 150.32 | -0.23 | 1 | *0.11* |
| | C vs. Adults | | -1.60 [-6.57-2.64] | 1.69 | 149.38 | -0.94 | 1 | *0.46* |
| | Ado vs. Adults | | -1.18 [-6.40-3.37] | 1.79 | 149.62 | -0.65 | 1 | *0.36* |
| | YC vs. C | High | -10.87 [-16.11-(-5.48)] | 1.94 | 150.29 | -5.58 | **<0.001** | *2.53* |
| | YC vs. Ado | | -14.52 [-20.03-(-8.91)] | 2.03 | 150.31 | -7.12 | **<0.001** | *3.40* |
| | YC vs. Adults | | -16.47 [-22.08-(-11.45)] | 1.95 | 149.89 | -8.44 | **<0.001** | *3.93* |
| | C vs. Ado | | -3.64 [-8.56-1.20] | 1.78 | 150.32 | -2.03 | 0.08 | *0.86* |
| | C vs. Adults | | -5.60 [-10.57-(-1.37)] | 1.69 | 149.38 | -3.30 | **<0.01** | *1.40* |
| | Ado vs. Adults | | -1.95 [-7.17-2.59] | 1.79 | 149.62 | -1.09 | 0.27 | *0.54* |

**Table S3 (Continued)**

| Variable | Comparison | Surface | Estimate [95%CI] | SE | df | t | *p-value** | *d* |
|---|---|---|---|---|---|---|---|---|
| **Variability** | | | | | | | | |
| GVI (au) | YC vs. C | Even | -6.57 [-12.71-(-0.56)] | 2.26 | 176.20 | -2.90 | **<0.01** | *1.20* |
| | YC vs. Ado | | -22.22 [-28.62-(-15.91)] | 2.36 | 176.22 | -9.39 | **<0.001** | *4.02* |
| | YC vs. Adults | | -25.63 [-31.49-(-19.37)] | 2.26 | 175.90 | -11.31 | **<0.001** | *4.58* |
| | C vs. Ado | | -15.65 [-21.21-(-10.05)] | 2.07 | 176.23 | -7.53 | **<0.001** | *2.82* |
| | C vs. Adults | | -19.05 [-24.04-(-13.52)] | 1.96 | 175.49 | -9.69 | **<0.001** | *3.39* |
| | Ado vs. Adults | | -3.40 [-8.73-2.43] | 2.08 | 175.68 | -1.63 | 0.10 | *0.57* |
| | YC vs. C | Medium | -4.41 [-10.55-1.60] | 2.26 | 176.20 | -1.95 | 0.11 | *0.81* |
| | YC vs. Ado | | -12.73 [-19.13-(-6.42)] | 2.36 | 176.22 | -5.38 | **<0.001** | *2.31* |
| | YC vs. Adults | | -13.15 [-19.01-(-6.86] | 2.26 | 175.90 | -5.80 | **<0.001** | *2.33* |
| | C vs. Ado | | -8.32 [-13.88-(-2.72)] | 2.07 | 176.23 | -4.00 | **<0.001** | *1.50* |
| | C vs. Adults | | -8.73 [-13.72-(-3.19)] | 1.96 | 175.49 | -4.44 | **<0.001** | *1.53* |
| | Ado vs. Adults | | -0.41 [-5.74-5.43] | 2.08 | 175.68 | -0.19 | 0.84 | *0.03* |
| | YC vs. C | High | -4.93 [-11.08-1.08] | 2.26 | 176.20 | -2.18 | 0.09 | *0.90* |
| | YC vs. Ado | | -8.21 [-14.61-(-1.90)] | 2.36 | 176.22 | -3.47 | **<0.01** | *1.49* |
| | YC vs. Adults | | -10.96 [-16.83-(-4.67)] | 2.26 | 175.90 | -4.84 | **<0.001** | *1.94* |
| | C vs. Ado | | -3.27 [-8.83-2.33] | 2.07 | 176.23 | -1.57 | 0.23 | *0.59* |
| | C vs. Adults | | -6.02 [-11.01-(-0.48)] | 1.96 | 175.49 | -3.06 | **<0.01** | *1.04* |
| | Ado vs. Adults | | -2.75 [-8.08-3.09] | 2.08 | 175.68 | -1.32 | 0.23 | *0.45* |

**Table S3 (Continued)**

| Variable | Comparison | Surface | Estimate [95%CI] | SE | df | t | *p-value** | *d* |
|---|---|---|---|---|---|---|---|---|
| **Asymmetry** | | | | | | | | |
| **SI stride length (%)** | YC vs. C | Even | 0.42 [-0.75-1.62] | 0.44 | 175.40 | 0.96 | 1 | *0.40* |
| | YC vs. Ado | | 0.39 [-0.84-1.64] | 0.46 | 175.42 | 0.85 | 1 | *0.37* |
| | YC vs. Adults | | 0.43 [-0.77-1.61] | 0.44 | 175.09 | 0.98 | 1 | *0.39* |
| | C vs. Ado | | -0.03 [-1.12-1.06] | 0.40 | 175.42 | -0.07 | 1 | *0.03* |
| | C vs. Adults | | 0.01 [-1.04-1.01] | 0.38 | 174.68 | 0.02 | 1 | *0.01* |
| | Ado vs. Adults | | 0.03 [-1.07-1.11] | 0.41 | 174.87 | 0.09 | 1 | *0.02* |
| | YC vs. C | Medium | 0.58 [-0.60-1.77] | 0.44 | 175.40 | 1.30 | 0.97 | *0.54* |
| | YC vs. Ado | | 0.71 [-0.52-1.96] | 0.46 | 175.42 | 1.53 | 0.75 | *0.66* |
| | YC vs. Adults | | 0.57 [0.63-1.75] | 0.44 | 175.09 | 1.29 | 0.97 | *0.51* |
| | C vs. Ado | | 0.13 [-0.95-1.22] | 0.40 | 175.42 | 0.33 | 1 | *0.12* |
| | C vs. Adults | | -0.01 [-1.05-1.00] | 0.38 | 174.68 | -0.01 | 1 | *0.02* |
| | Ado vs. Adults | | -0.14 [-1.25-0.93] | 0.41 | 174.87 | -0.34 | 1 | *0.15* |
| | YC vs. C | High | 3.10 [1.93-4.30] | 0.44 | 175.40 | 6.97 | **<0.001** | *2.86* |
| | YC vs. Ado | | 2.98 [1.75-4.23] | 0.46 | 175.42 | 6.41 | **<0.001** | *2.75* |
| | YC vs. Adults | | 3.14 [1.94-4.31] | 0.44 | 175.09 | 7.03 | **<0.001** | *2.87* |
| | C vs. Ado | | -0.11 [-1.21-0.97] | 0.40 | 175.42 | -0.29 | 1 | *0.11* |
| | C vs. Adults | | 0.03 [-1.02-1.04] | 0.38 | 174.68 | 0.08 | 1 | *0.01* |
| | Ado vs. Adults | | 0.15 [-0.96-1.22] | 0.41 | 174.87 | 0.37 | 1 | *0.12* |

***Notes:*** **p-values were adjusted using the Holm-Bonferroni correction. Estimates represent the mean difference between age groups for a surface with 95% Confidence Intervals (95% CI). Cohen's d was calculated using the model's residual standard deviation as the denominator.* ***Abbreviations:*** *adolescents (Ado); anteroposterior (AP); arbitrary unit (au); children (C); Cohen's d (d); degrees of freedom (df); gait variability index (GVI); leg length (L0); millimeter* *(mm); margin of stability (MoS); standard error (SE); symmetry index (SI); spectral arc length (SPARC) ; young children (YC).*

**Table S4:** Post-hoc pairwise comparisons between surface by age for gait variables showing a significant effect of the interaction Age × Surface.

| Variable | Comparison | Age | Estimate [95%CI] | SE | df | t | *p-value** | *d* |
|---|---|---|---|---|---|---|---|---|
| **Pace** | | | | | | | | |
| **Normalized step length (au)** | Even *vs.* Medium | YC | 0.04 [0.00-0.08] | 0.02 | 128 | 2.71 | **<0.001** | *1.10* |
| | Even *vs.* High | | 0.09 [0.05-0.12] | 0.02 | 128 | 5.69 | **<0.001** | *2.32* |
| | Medium *vs.* High | | 0.05 [0.01-0.08] | 0.02 | 128 | 2.97 | **<0.001** | *1.21* |
| | Even *vs.* Medium | C | 0.01 [-0.02-0.04] | 0.01 | 128 | 0.53 | 0.69 | *0.17* |
| | Even *vs.* High | | 0.02 [-0.01-0.05] | 0.01 | 128 | 1.48 | 0.42 | *0.47* |
| | Medium *vs.* High | | 0.01 [-0.01-0.04] | 0.01 | 128 | 0.95 | 0.69 | *0.30* |
| | Even *vs.* Medium | Ado | -0.01 [-0.04-0.02] | 0.01 | 128 | -0.88 | 0.38 | *0.31* |
| | Even *vs.* High | | 0.02 [-0.01-0.06] | 0.01 | 128 | 1.75 | 0.17 | *0.62* |
| | Medium *vs.* High | | 0.03 [-0.00-0.07] | 0.01 | 128 | 2.62 | **0.03** | *0.93* |
| | Even *vs.* Medium | Adults | -0.01 [-0.04-0.02] | 0.01 | 128 | -0.75 | 0.46 | *0.24* |
| | Even *vs.* High | | 0.02 [-0.01-0.04] | 0.01 | 128 | 1.36 | 0.35 | *0.42* |
| | Medium *vs.* High | | 0.03 [-0.01-0.05] | 0.01 | 128 | 2.10 | 0.11 | *0.67* |

**Table S4 (Continued)**

| Variable | Comparison | Age | Estimate [95%CI] | SE | df | t | *p-value** | *d* |
|---|---|---|---|---|---|---|---|---|
| | | | **Rythm** | | | | | |
| **SPARC (au)** | Even *vs.* Medium | YC | 1.64 [0.47-2.82] | 0.49 | 128 | 3.38 | **<0.001** | *1.38* |
| | Even *vs.* High | | 3.25 [2.07-4.43] | 0.49 | 128 | 6.69 | **<0.001** | *2.73* |
| | Medium *vs.* High | | 1.61 [0.42-2.78] | 0.49 | 128 | 3.30 | **<0.001** | *1.35* |
| | Even *vs.* Medium | C | 0.15 [-0.76-1.06] | 0.38 | 128 | 0.40 | 0.70 | *0.13* |
| | Even *vs.* High | | 0.83 [-0.09-1.74] | 0.38 | 128 | 2.20 | 0.09 | *0.70* |
| | Medium *vs.* High | | 0.68 [-0.24-1.59] | 0.38 | 128 | 1.80 | 0.15 | *0.57* |
| | Even *vs.* Medium | Ado | 0.96 [-0.06-1.98] | 0.42 | 128 | 2.29 | 0.07 | *0.81* |
| | Even *vs.* High | | 0.80 [-0.22-1.82] | 0.42 | 128 | 1.90 | 0.12 | *0.67* |
| | Medium *vs.* High | | -0.16 [-1.18-0.85] | 0.42 | 128 | -0.39 | 0.70 | *0.14* |
| | Even *vs.* Medium | Adults | 0.89 [-0.02-1.80] | 0.38 | 128 | 2.37 | 0.06 | *0.75* |
| | Even *vs.* High | | 0.87 [-0.04-1.78] | 0.38 | 128 | 2.31 | 0.06 | *0.73* |
| | Medium *vs.* High | | -0.02 [-0.94-0.89] | 0.38 | 128 | -0.06 | 0.95 | *0.02* |

**Table S4 (Continued)**

| Variable | Comparison | Age | Estimate [95%CI] | SE | df | t | *p-value** | *d* |
|---|---|---|---|---|---|---|---|---|
| **Dynamic stability** | | | | | | | | |
| **Normalized step width (au)** | Even *vs*. Medium | YC | -0.00 [-0.02-0.02] | 0.01 | 128 | -0.44 | 0.65 | *0.18* |
| | Even *vs*. High | | -0.03 [-0.05-(-0.01)] | 0.01 | 128 | -3.80 | **<0.001** | *1.55* |
| | Medium *vs*. High | | -0.03 [-0.05-(-0.01] | 0.01 | 128 | -3.36 | **<0.001** | *1.37* |
| | Even *vs*. Medium | C | -0.00 [-0.02-0.01] | 0.01 | 128 | -0.22 | 0.83 | *0.07* |
| | Even *vs*. High | | 0.00 [-0.01-0.02] | 0.01 | 128 | 1.16 | 0.51 | *0.36* |
| | Medium *vs*. High | | 0.00 [-0.01-0.02] | 0.01 | 128 | 1.38 | 0.51 | *0.44* |
| | Even *vs*. Medium | Ado | -0.00 [-0.01-0.02] | 0.01 | 128 | 0.63 | 0.53 | *0.22* |
| | Even *vs*. High | | -0.01 [-0.03-0.01] | 0.01 | 128 | -1.32 | 0.38 | *0.47* |
| | Medium *vs*. High | | -0.01 [-0.03-0.00] | 0.01 | 128 | -1.94 | 0.16 | *0.69* |
| | Even *vs*. Medium | Adults | -0.00 [-0.02-0.01] | 0.01 | 128 | -1.28 | 0.41 | *0.40* |
| | Even *vs*. High | | -0.02 [-0.03-0.00] | 0.01 | 128 | -2.49 | **0.04** | *0.78* |
| | Medium *vs*. High | | -0.00 [-0.02-0.01] | 0.01 | 128 | -1.20 | 0.41 | *0.38* |

#### Table S4 (Continued)

| Variable | Comparison | Age | Estimate [95%CI] | SE | df | t | *p-value** | *d* |
|---|---|---|---|---|---|---|---|---|
| **Dynamic stability** | | | | | | | | |
| **MoS AP (mm)** | Even *vs.* Medium | YC | 14.98 [-11.64-41.60] | 10.97 | 128 | 1.36 | 0.17 | *0.56* |
| | Even *vs.* High | | 67.54 [40.91-94.16] | 10.97 | 128 | 6.15 | **<0.001** | *2.51* |
| | Medium *vs.* High | | 52.56 [25.93-79.18] | 10.97 | 128 | 4.79 | **<0.001** | *1.96* |
| | Even *vs.* Medium | C | 0.78 [-19.84-21.41] | 8.50 | 128 | 0.09 | 0.92 | *0.03* |
| | Even *vs.* High | | 28.98 [8.36-49.60] | 8.50 | 128 | 3.41 | **<0.001** | *1.07* |
| | Medium *vs.* High | | 28.19 [7.57-48.81] | 8.50 | 128 | 3.32 | **<0.001** | *1.05* |
| | Even *vs.* Medium | Ado | 17.55 [-5.50-40.61] | 9.50 | 128 | 1.85 | 0.13 | *0.65* |
| | Even *vs.* High | | 26.43 [3.38-49.49] | 9.50 | 128 | 2.78 | **0.02** | *0.98* |
| | Medium *vs.* High | | 8.88 [-14.17-31.94] | 9.50 | 128 | 0.93 | 0.35 | *0.33* |
| | Even *vs.* Medium | Adults | 10.98 [-9.64-31.60] | 8.50 | 128 | 1.29 | 0.40 | *0.41* |
| | Even *vs.* High | | 13.24 [-7.38-33.86] | 8.50 | 128 | 1.56 | 0.37 | *0.49* |
| | Medium *vs.* High | | 2.26 [-18.35-22.88] | 8.50 | 128 | 0.27 | 0.79 | *0.08* |

#### Table S4 (Continued)

| Variable | Comparison | Age | Estimate [95%CI] | SE | df | t | *p-value** | *d* |
|---|---|---|---|---|---|---|---|---|
| **Dynamic stability** | | | | | | | | |
| **Normalized MoS AP (%L0)** | Even *vs.* Medium | YC | 3.19 [-1.03-7.41] | 1.74 | 128 | 1.83 | 0.07 | *0.75* |
| | Even *vs.* High | | 13.57 [9.36-17.80] | 1.74 | 128 | 7.80 | **<0.001** | *3.18* |
| | Medium *vs.* High | | 10.39 [6.17-14.61] | 1.74 | 128 | 5.97 | **<0.001** | *2.44* |
| | Even *vs.* Medium | C | 0.11 [-3.16-3.38] | 1.35 | 128 | 0.08 | 0.93 | *0.03* |
| | Even *vs.* High | | 4.37 [1.10-7.64] | 1.35 | 128 | 3.24 | **<0.001** | *1.02* |
| | Medium *vs.* High | | 4.25 [0.99-7.52] | 1.35 | 128 | 3.16 | **<0.001** | *0.99* |
| | Even *vs.* Medium | Ado | 2.01 [-1.64-5.67] | 1.51 | 128 | 1.34 | 0.37 | *0.47* |
| | Even *vs.* High | | 3.04 [-0.61-6.70] | 1.51 | 128 | 2.01 | 0.14 | *0.71* |
| | Medium *vs.* High | | 1.03 [-2.63-4.68] | 1.51 | 128 | 0.68 | 0.49 | *0.24* |
| | Even *vs.* Medium | Adults | 1.21 [-2.06-4.48] | 1.35 | 128 | 0.90 | 0.84 | *0.28* |
| | Even *vs.* High | | 1.46 [-1.81-4.73] | 1.35 | 128 | 1.08 | 0.84 | *0.34* |
| | Medium *vs.* High | | 0.25 [-3.02-3.52] | 1.35 | 128 | 0.19 | 0.85 | *0.06* |

#### Table S4 (Continued)

| **Variable** | **Comparison** | **Age** | **Estimate [95%CI]** | **SE** | **df** | **t** | ***p-value**** | ***d*** |
|---|---|---|---|---|---|---|---|---|
| | | | **Variability** | | | | | |
| **GVI (au)** | Even *vs.* Medium | YC | 2.06 [-3.43-7.55] | 2.26 | 128 | 0.91 | 0.37 | *0.37* |
| | Even *vs.* High | | 6.83 [1.33-12.32] | 2.26 | 128 | 3.01 | **<0.001** | *1.23* |
| | Medium *vs.* High | | 4.77 [-0.72-10.26] | 2.26 | 128 | 2.11 | 0.07 | *0.76* |
| | Even *vs.* Medium | C | 4.21 [-0.04-8.47] | 1.75 | 128 | 2.40 | **0.03** | *0.86* |
| | Even *vs.* High | | 8.46 [4.21-12.71] | 1.75 | 128 | 4.83 | **<0.001** | *1.53* |
| | Medium *vs.* High | | 4.25 [-0.00-8.50] | 1.75 | 128 | 2.42 | **0.03** | *0.86* |
| | Even *vs.* Medium | Ado | 11.54 [6.79-16.30] | 1.96 | 128 | 5.89 | **<0.001** | *2.08* |
| | Even *vs.* High | | 20.84 [16.08-25.59] | 1.96 | 128 | 10.63 | **<0.001** | *3.76* |
| | Medium *vs.* High | | 9.30 [4.54-14.05] | 1.96 | 128 | 4.74 | **<0.001** | *1.68* |
| | Even *vs.* Medium | Adults | 14.54 [10.28-18.79] | 1.75 | 128 | 8.29 | **<0.001** | *2.62* |
| | Even *vs.* High | | 21.49 [17.24-25.75] | 1.75 | 128 | 12.26 | **<0.001** | *3.88* |
| | Medium *vs.* High | | 6.96 [2.71-11.21] | 1.75 | 128 | 3.97 | **<0.001** | *1.26* |

**Table S4 (Continued)**

| Variable | Comparison | Age | Estimate [95%CI] | SE | df | t | *p-value** | *d* |
|---|---|---|---|---|---|---|---|---|
| **Asymmetry** | | | | | | | | |
| **SI stride length (%)** | Even *vs.* Medium | YC | -0.34 [-1.42-0.74] | 0.44 | 128 | -0.76 | 0.44 | *0.31* |
| | Even *vs.* High | | -3.01 [-4.09-(-1.93)] | 0.44 | 128 | -6.77 | **<0.001** | *2.76* |
| | Medium *vs.* High | | -2.67 [-3.75-(-1.59)] | 0.44 | 128 | -6.00 | **<0.001** | *2.45* |
| | Even *vs.* Medium | C | -0.19 [-1.02-0.65] | 0.34 | 128 | -0.55 | 1 | *0.17* |
| | Even *vs.* High | | -0.33 [-1.17-0.50] | 0.34 | 128 | -0.96 | 1 | *0.30* |
| | Medium *vs.* High | | -0.14 [-0.98-0.69] | 0.34 | 128 | -0.41 | 1 | *0.13* |
| | Even *vs.* Medium | Ado | -0.02 [-0.95-0.91] | 0.38 | 128 | -0.05 | 0.95 | *0.02* |
| | Even *vs.* High | | -0.42 [-1.35-0.51] | 0.38 | 128 | -1.09 | 0.83 | *0.38* |
| | Medium *vs.* High | | -0.40 [-1.33-0.53] | 0.38 | 128 | -1.04 | 0.83 | *0.37* |
| | Even *vs.* Medium | Adults | -0.20 [-1.03-0.64] | 0.34 | 128 | -0.58 | 1 | *0.18* |
| | Even *vs.* High | | -0.30 [-1.14-0.53] | 0.34 | 128 | -0.89 | 1 | *0.28* |
| | Medium *vs.* High | | -0.10 [-0.94-0.73] | 0.34 | 128 | -0.31 | 1 | *0.10* |

***Notes:*** **p-values were adjusted using the Holm-Bonferroni correction. Estimates represent the mean difference between surfaces for an age group with 95% Confidence Intervals (95% CI). Cohen's d was calculated using the model's residual standard deviation as the denominator.* ***Abbreviations:*** *adolescents (Ado); anteroposterior (AP); arbitrary unit (au); children (C); Cohen's d (d); degrees of freedom (df); gait variability index (GVI); leg length (L0); millimeter (mm); margin of stability (MoS); standard error (SE); symmetry index (SI); spectral arc length (SPARC); young children (YC).*

**Table S5:** Post-hoc pairwise comparisons between each age group and adults for gait variables showing a significant effect of Age.

| Variable | Comparison | Estimate [95%CI] | SE | df | t | *p-value** | *d* |
|---|---|---|---|---|---|---|---|
| **Pace** | | | | | | | |
| **Gait Speed (m/s)** | YC vs. Adults | -0.30 [-0.46-(-0.14)] | 0.06 | 64 | -5.17 | **<0.001** | *3.53* |
| | C vs. Adults | -0.11 [-0.25--0.03] | 0.05 | 64 | -2.19 | 0.10 | *1.29* |
| | Ado vs. Adults | -0.04 [-0.18-0.11] | 0.05 | 64 | -0.68 | 0.50 | *0.43* |
| **Step length (m)** | YC vs. Adults | -0.28 [-0.34-(-0.21)] | 0.02 | 64 | -11.70 | **<0.001** | *10.39* |
| | C vs. Adults | -0.14 [-0.19-(-0.08)] | 0.02 | 64 | -6.67 | **<0.001** | *5.13* |
| | Ado vs. Adults | -0.03 [-0.09-0.03] | 0.02 | 64 | -1.46 | 0.14 | *1.19* |
| **Norm step length (au)** | YC vs. Adults | 0.08 [0.01-0.16] | 0.03 | 64 | 3.05 | **0.01** | *2.15* |
| | C vs. Adults | 0.08 [0.02-0.14] | 0.02 | 64 | 3.39 | **<0.001** | *2.07* |
| | Ado vs. Adults | -0.00 [-0.07-0.06] | 0.02 | 64 | -0.13 | 1.00 | *0.09* |
| **Stride length (m)** | YC vs. Adults | -0.55 [-0.68-(-0.42)] | 0.04 | 64 | -11.62 | **<0.001** | *10.36* |
| | C vs. Adults | -0.27 [-0.38-(-0.16)] | 0.04 | 64 | -6.64 | **<0.001** | *5.12* |
| | Ado vs. Adults | -0.06 [-0.18-0.06] | 0.04 | 64 | -1.40 | 0.17 | *1.14* |
| **Walk ratio (cm.min.pas$^{-1}$)** | YC vs. Adults | -0.33 [-0.40-(-0.26)] | 0.02 | 64 | -13.37 | **<0.001** | *14.60* |
| | C vs. Adults | -0.18 [-0.24-(-0.12)] | 0.02 | 64 | -8.53 | **<0.001** | *8.07* |
| | Ado vs. Adults | -0.04 [-0.10-0.03] | 0.02 | 64 | -1.62 | 0.11 | *1.63* |
| **Norm Walk ratio (au)** | YC vs. Adults | 0.29 [0.16-0.41] | 0.05 | 64 | 6.34 | **<0.001** | *3.67* |
| | C vs. Adults | 0.20 [0.09-0.31] | 0.04 | 64 | 5.08 | **<0.001** | *2.55* |
| | Ado vs. Adults | 0.00 [-0.11-0.11] | 0.04 | 64 | 0.01 | 0.99 | *0.01* |

**Table S5 (Continued)**

| Variable | Comparison | Estimate [95%CI] | SE | df | t | *p-value** | *d* |
|---|---|---|---|---|---|---|---|
| **Rythm** | | | | | | | |
| **Double support time (%)** | YC vs. Adults | 1.59 [-0.92-4.10] | 0.92 | 64 | 1.73 | 0.36 | *1.11* |
| | C vs. Adults | -1.29 [-3.46-0.88] | 0.80 | 64 | -1.61 | 0.56 | *0.90* |
| | Ado vs. Adults | 0.38 [-1.93-2.69] | 0.85 | 64 | 0.45 | 0.66 | *0.27* |
| **Single support time (%)** | YC vs. Adults | -0.78 [-2.72-(-0.24)] | 0.46 | 64 | -1.72 | 0.33 | *1.00* |
| | C vs. Adults | 0.69 [-0.38-1.77] | 0.39 | 64 | 1.76 | 0.33 | *0.89* |
| | Ado vs. Adults | -0.19 [-1.32-0.95] | 0.42 | 64 | -0.46 | 0.65 | *0.24* |
| **Stance time (s)** | YC vs. Adults | -0.12 [-0.18-(-0.06)] | 0.02 | 64 | -5.50 | **<0.001** | *3.42* |
| | C vs. Adults | -0.08 [-0.13-(-0.03)] | 0.02 | 64 | -4.33 | **<0.001** | *2.33* |
| | Ado vs. Adults | -0.01 [-0.06-0.05] | 0.02 | 64 | -0.30 | 0.76 | *0.17* |
| **Swing time (s)** | YC vs. Adults | -0.08 [-0.11-(-0.05)] | 0.01 | 64 | -8.18 | **<0.001** | *5.01* |
| | C vs. Adults | -0.04 [-0.06-(-0.01)] | 0.01 | 64 | -4.33 | **<0.001** | *1.87* |
| | Ado vs. Adults | -0.01 [-0.03-0.02] | 0.01 | 64 | -0.74 | 0.46 | *0.42* |
| **Step time (s)** | YC vs. Adults | -0.10 [-0.14-(-0.06)] | 0.01 | 64 | -6.86 | **<0.001** | *4.37* |
| | C vs. Adults | -0.06 [-0.09-(-0.02)] | 0.01 | 64 | -4.64 | **<0.001** | *2.56* |
| | Ado vs. Adults | -0.01 [-0.04-0.03] | 0.01 | 64 | -0.48 | 0.63 | *0.28* |
| **Stride time (s)** | YC vs. Adults | -0.20 [-0.28-(-0.12)] | 0.03 | 64 | -6.64 | **<0.001** | *4.13* |
| | C vs. Adults | -0.12 [-0.19-(-0.05)] | 0.03 | 64 | -4.54 | **<0.001** | *2.44* |
| | Ado vs. Adults | -0.01 [-0.09-0.06] | 0.03 | 64 | -0.46 | 0.65 | *0.26* |
| **Cadence (step/min)** | YC vs. Adults | 25.76 [16.70-34.82] | 3.33 | 64 | 7.74 | **<0.001** | *4.93* |
| | C vs. Adults | 14.04 [6.19-21.88] | 2.88 | 64 | 4.87 | **<0.001** | *2.69* |
| | Ado vs. Adults | 1.58 [-6.75-9.90] | 3.06 | 64 | 0.52 | 0.61 | *0.30* |

**Table S5 (Continued)**

| Variable | Comparison | Estimate [95%CI] | SE | df | t | *p-value** | *d* |
|---|---|---|---|---|---|---|---|
| **Norm cadence (au)** | YC vs. Adults | -0.04 [-0.08-(-0.01)] | 0.01 | 64 | -3.13 | **0.02** | *1.84* |
| | C vs. Adults | -0.02 [-0.05-0.01] | 0.01 | 64 | -1.54 | 0.38 | *0.79* |
| | Ado vs. Adults | -0.00 [-0.04-0.03] | 0.01 | 64 | -0.29 | 0.77 | *0.16* |
| **SPARC (au)** | YC vs. Adults | -3.15 [-4.10-(-2.19)] | 0.35 | 64 | -8.98 | **<0.001** | *2.65* |
| | C vs. Adults | -0.89 [-1.72-(-0.06)] | 0.30 | 64 | -2.93 | **0.01** | *0.75* |
| | Ado vs. Adults | -0.56 [-1.44-0.32] | 0.32 | 64 | -1.74 | 0.17 | *0.47* |
| **Dynamic stability** | | | | | | | |
| **Norm step width (au)** | YC vs. Adults | 0.06 [0.03-0.10] | 0.01 | 64 | 5.37 | **<0.001** | *3.24* |
| | C vs. Adults | 0.03 [0.00-0.05] | 0.01 | 64 | 2.30 | 0.05 | *1.28* |
| | Ado vs. Adults | 0.01 [-0.03-0.03] | 0.01 | 64 | 0.00 | 0.99 | *0.00* |
| **MoS A.P. (mm)** | YC vs. Adults | 24.67 [-8.08-57.41] | 12.03 | 64 | 2.06 | 0.18 | *0.92* |
| | C vs. Adults | 18.44 [-9.92-46.80] | 10.42 | 64 | 1.77 | 0.24 | *0.69* |
| | Ado vs. Adults | -5.70 [-35.77-24.39] | 11.05 | 64 | -0.52 | 1.00 | *0.21* |
| **MoS M.L (mm)** | YC vs. Adults | -15.22 [-25.22-(-5.23)] | 3.67 | 64 | -4.15 | **<0.001** | *2.42* |
| | C vs. Adults | -14.91 [-23.56-(-6.25)] | 3.18 | 64 | -4.69 | **<0.001** | *2.36* |
| | Ado vs. Adults | -2.43 [-11.61-6.75] | 3.37 | 64 | -0.72 | 0.95 | *0.38* |
| **Norm MoS A.P (%L0)** | YC vs. Adults | -9.34 [-13.50-(-5.19)] | 1.53 | 64 | -6.11 | **<0.001** | *2.19* |
| | C vs. Adults | -3.66 [-7.27-(-0.06)] | 1.32 | 64 | -2.77 | **0.02** | *0.86* |
| | Ado vs. Adults | -1.51 [-5.33-2.31] | 1.40 | 64 | -1.07 | 0.29 | *0.35* |
| **Norm MoS M.L (%L0)** | YC vs. Adults | 4.20 [2.77-5.64] | 0.53 | 64 | 8.00 | **<0.001** | *4.80* |
| | C vs. Adults | 1.08 [-0.16-2.32] | 0.46 | 64 | 2.37 | 0.06 | *1.23* |
| | Ado vs. Adults | 0.05 [-1.26-1.37] | 0.48 | 64 | 0.11 | 0.91 | *0.06* |

#### Table S5 (Continued)

| Variable | Comparison | Estimate [95%CI] | SE | df | t | *p-value** | *d* |
|---|---|---|---|---|---|---|---|
| **Variability** | | | | | | | |
| **GVI (au)** | YC vs. Adults | -16.36 [-20.63-(-12.10)] | 1.57 | 64 | -10.44 | **<0.001** | *2.95* |
| | C vs. Adults | -11.00 [-14.69-(-7.30)] | 1.36 | 64 | -8.10 | **<0.001** | *1.98* |
| | Ado vs. Adults | -1.93 [-5.85-1.99] | 1.44 | 64 | -1.34 | 0.18 | *0.34* |
| **CV gait speed (%)** | YC vs. Adults | 8.77 [6.35-11.19] | 0.89 | 64 | 9.86 | **<0.001** | *3.93* |
| | C vs. Adults | 3.72 [1.63-5.82] | 0.77 | 64 | 4.84 | **<0.001** | *1.67* |
| | Ado vs. Adults | 0.10 [-2.12-2.32] | 0.82 | 64 | 0.12 | 0.90 | *0.05* |
| **CV step width (%)** | YC vs. Adults | 19.65 [9.85-29.44] | 3.60 | 64 | 5.46 | **<0.001** | *2.33* |
| | C vs. Adults | 17.31 [8.83-25.79] | 3.11 | 64 | 5.56 | **<0.001** | *2.05* |
| | Ado vs. Adults | 4.21 [-4.78-13.21] | 3.30 | 64 | 1.27 | 0.41 | *0.50* |
| **Asymmetry** | | | | | | | |
| **SI stride length (%)** | YC vs. Adults | 1.37 [0.54-2.20] | 0.30 | 64 | 4.50 | **<0.001** | *1.26* |
| | C vs. Adults | -0.01 [-0.73-0.71] | 0.27 | 64 | -0.04 | 1.00 | *0.01* |
| | Ado vs. Adults | -0.03 [-0.76-0.76] | 0.28 | 64 | -0.01 | 1.00 | *0.00* |
| **SI double support time (%)** | YC vs. Adults | 1.51 [0.46-2.56] | 0.38 | 64 | 3.91 | **<0.001** | *0.82* |
| | C vs. Adults | 0.18 [-0.73-1.09] | 0.33 | 64 | 0.54 | 1.00 | *0.10* |
| | Ado vs. Adults | 0.42 [-0.55-1.38] | 0.35 | 64 | 1.18 | 0.73 | *0.23* |
| **SI step width (%)** | YC vs. Adults | 15.10 [6.39-23.80] | 3.20 | 64 | 4.72 | **<0.001** | *1.02* |
| | C vs. Adults | 4.42 [-3.12-11.96] | 2.77 | 64 | 1.60 | 0.35 | *0.30* |
| | Ado vs. Adults | 1.85 [-6.15-9.84] | 2.94 | 64 | 0.63 | 0.77 | *0.13* |

***Notes:*** **p-values were adjusted using the Holm-Bonferroni correction. Estimates represent the mean difference between age groups with 95% Confidence Intervals (95% CI). Cohen's d was calculated using the model's residual standard deviation as the denominator.* ***Abbreviations:*** *adolescents (Ado); anteroposterior (AP); arbitrary unit (au); children (C); centimeter (cm); coefficient of variation (CV); Cohen's d (d); degrees of freedom (df); gait variability index (GVI); leg length (L0); meter (m); minute (min); mediolateral (ML); millimeter (mm); margin of stability (MoS); second (s); standard error (SE); symmetry index (SI); spectral arc length (SPARC); young children (YC).*

**Table S6:** Post-hoc pairwise comparisons between each irregular surface (Medium, High) and even for gait variables showing a significant effect of Surface.

| Variable | Comparison | Estimate [95%CI] | SE | df | t | *p-value** | *d* |
|---|---|---|---|---|---|---|---|
| **Pace** | | | | | | | |
| **Gait Speed (m/s)** | E vs. Medium | 0.04 [0.01-0.08] | 0.01 | 128 | 2.83 | **<0.001** | *0.50* |
| | E vs. High | 0.11 [0.07-0.15] | 0.01 | 128 | 7.38 | **<0.001** | *1.29* |
| **Norm gait speed (au)** | E vs. Medium | 0.02 [0.00-0.03] | 0.01 | 128 | 2.98 | **<0.001** | *0.52* |
| | E vs. High | 0.04 [0.03-0.06] | 0.01 | 128 | 7.35 | **<0.001** | *1.29* |
| **Step length (m)** | E vs. Medium | 0.00 [-0.01-0.01] | 0.00 | 128 | 0.34 | 0.73 | *0.06* |
| | E vs. High | 0.02 [0.01-0.03] | 0.00 | 128 | 4.90 | **<0.001** | *0.86* |
| **Norm step length (au)** | E vs. Medium | 0.00 [-0.01-0.02] | 0.00 | 128 | 1.04 | 0.30 | *0.18* |
| | E vs. High | 0.04 [0.02-0.05] | 0.01 | 128 | 5.47 | **<0.001** | *0.96* |
| **Stride length (m)** | E vs. Medium | 0.00 [-0.02-0.03] | 0.01 | 128 | 0.35 | 0.73 | *0.06* |
| | E vs. High | 0.04 [0.02-0.07] | 0.01 | 128 | 4.36 | **<0.001** | *0.83* |
| **Walk ratio (cm.min.pas$^{-1}$)** | E vs. Medium | -0.02 [-0.03-(-0.01)] | 0.00 | 128 | -4.02 | **<0.001** | *0.70* |
| | E vs. High | -0.01 [-0.02-(-0.00)] | 0.00 | 128 | -3.01 | **<0.001** | *0.53* |
| **Norm Walk ratio (au)** | E vs. Medium | -0.03 [-0.07-(-0.00)] | 0.01 | 128 | -2.56 | **0.03** | *0.45* |
| | E vs. High | -0.02 [-0.05-0.01] | 0.01 | 128 | -1.37 | 0.35 | *0.24* |
| **Rythm** | | | | | | | |
| **Double support time (%)** | E vs. Medium | 1.45 [0.84-2.06] | 0.25 | 128 | 5.79 | **<0.001** | *1.01* |
| | E vs. High | 1.76 [1.15-2.37] | 0.25 | 128 | 7.03 | **<0.001** | *1.23* |
| **Single support time (%)** | E vs. Medium | -0.68 [-1.01-(-0.34)] | 0.14 | 128 | -4.94 | **<0.001** | *0.87* |
| | E vs. High | -0.75 [-1.08-(-0.42)] | 0.14 | 128 | -5.46 | **<0.001** | *0.96* |
| **Stance time (s)** | E vs. Medium | -0.01 [-0.03-0.00] | 0.01 | 128 | -2.01 | **0.05** | *0.35* |
| | E vs. High | -0.03 [-0.04-(-0.01)] | 0.01 | 128 | -4.51 | **<0.001** | *0.79* |

#### Table S6 (Continued)

| Variable | Comparison | Estimate [95%CI] | SE | df | t | *p-value** | *d* |
|---|---|---|---|---|---|---|---|
| **Swing time (s)** | E vs. Medium | -0.02 [-0.03-(-0.01)] | 0.00 | 128 | -7.01 | **<0.001** | *1.23* |
| | E vs. High | -0.03 [-0.04-(-0.02)] | 0.00 | 128 | -11.33 | **<0.001** | *1.99* |
| **Step time (s)** | E vs. Medium | -0.02 [-0.03-(-0.01)] | 0.00 | 128 | -3.79 | **<0.001** | *0.67* |
| | E vs. High | -0.03 [-0.04-(-0.02)] | 0.00 | 128 | -6.92 | **<0.001** | *1.21* |
| **Stride time (s)** | E vs. Medium | -0.03 [-0.05-(-0.01)] | 0.01 | 128 | -3.77 | **<0.001** | *0.66* |
| | E vs. High | -0.06 [-0.08-(-0.04)] | 0.01 | 128 | -7.00 | **<0.001** | *1.23* |
| **Cadence (step/min)** | E vs. Medium | 3.56 [1.34-5.78] | 0.92 | 128 | 3.89 | **<0.001** | *0.68* |
| | E vs. High | 6.10 [3.88-8.32] | 0.92 | 128 | 6.69 | **<0.001** | *1.17* |
| **Norm cadence (au)** | E vs. Medium | 0.02 [0.01-0.03] | 0.00 | 128 | 4.11 | **<0.001** | *0.72* |
| | E vs. High | 0.03 [0.02-0.04] | 0.00 | 128 | 6.88 | **<0.001** | *1.26* |
| **SPARC (au)** | E vs. Medium | 0.91 [0.41-1.42] | 0.21 | 128 | 4.37 | **<0.001** | *0.77* |
| | E vs. High | 1.44 [0.93-1.94] | 0.21 | 128 | 6.88 | **<0.001** | *1.21* |
| **Dynamic stability** | | | | | | | |
| **Step width (cm)** | E vs. Medium | -0.19 [-0.81-0.43] | 0.25 | 128 | -0.74 | 0.46 | *0.13* |
| | E vs. High | -0.87 [-1.49-(-0.25)] | 0.25 | 128 | -3.40 | **<0.001** | *0.60* |
| **Norm step width (au)** | E vs. Medium | -0.00 [-9.31-3.23] | 0.00 | 128 | -0.62 | 0.54 | *0.21* |
| | E vs. High | -0.01 [-5.71-6.83] | 0.00 | 128 | -3.47 | **<0.001** | *0.04* |
| **MoS A.P. (mm)** | E vs. Medium | 11.07 [-0.36-22.50] | 4.71 | 128 | 2.35 | **0.02** | *0.41* |
| | E vs. High | 34.05 [22.62-45.48] | 4.71 | 128 | 7.23 | **<0.001** | *1.27* |
| **Norm MoS A.P (%L0)** | E vs. Medium | 1.63 [-0.18-3.44] | 0.75 | 128 | 2.18 | 0.03 | *0.38* |
| | E vs. High | 5.61 [3.80-7.42] | 0.75 | 128 | 7.51 | **<0.001** | *1.32* |
| | E vs. High | -1.49 [-2.27-(-0.71)] | 0.32 | 128 | -4.63 | **<0.001** | *0.81* |

#### Table S6 (Continued)

| Variable | Comparison | Estimate [95%CI] | SE | df | t | *p-value** | *d* |
|---|---|---|---|---|---|---|---|
| **Variability** | | | | | | | |
| **GVI (au)** | E vs. Medium | 8.09 [5.73-10.44] | 0.97 | 128 | 8.32 | **<0.001** | *1.46* |
| | E vs. High | 14.41 [12.05-16.76] | 0.97 | 128 | 14.83 | **<0.001** | *2.60* |
| **CV gait speed (%)** | E vs. Medium | -0.84 [-1.79-0.11] | 0.39 | 128 | -2.14 | **0.03** | *0.38* |
| | E vs. High | -1.89 [-2.84-(-0.94)] | 0.39 | 128 | -4.84 | **<0.001** | *0.85* |
| **CV step width (%)** | E vs. Medium | -7.64 [-11.23-(-4.05)] | 1.48 | 128 | -5.16 | **<0.001** | *0.90* |
| | E vs. High | -10.42 [-14.01-(-6.83)] | 1.48 | 128 | -7.04 | **<0.001** | *1.23* |
| **Asymmetry** | | | | | | | |
| **SI stride length (%)** | E vs. Medium | -0.19 [-0.65-0.28] | 0.19 | 128 | -0.98 | 0.33 | *0.17* |
| | E vs. High | -1.02 [-1.48-(-0.55)] | 0.19 | 128 | -5.32 | **<0.001** | *0.93* |
| **SI double support time (%)** | E vs. Medium | -0.72 [-1.50-0.06] | 0.32 | 128 | -1.23 | **0.05** | *0.39* |
| | E vs. High | -1.49 [-2.27-(-0.71)] | 0.32 | 128 | -4.63 | **<0.001** | *0.81* |

***Notes:*** **p-values were adjusted using the Holm-Bonferroni correction. Estimates represent the mean difference between surfaces with 95% Confidence Intervals (95% CI). Cohen's d was calculated using the model's residual standard deviation as the denominator.* ***Abbreviations:*** *anteroposterior (AP); arbitrary unit (au); centimeter (cm); coefficient of variation (CV); Cohen's d (d); degrees of freedom (df); Even (E); gait variability index (GVI); leg length (L0); meter (m); minute (min); mediolateral (ML); millimeter (mm); margin of stability (MoS); second (s); standard error (SE); symmetry index (SI); spectral arc length (SPARC).*

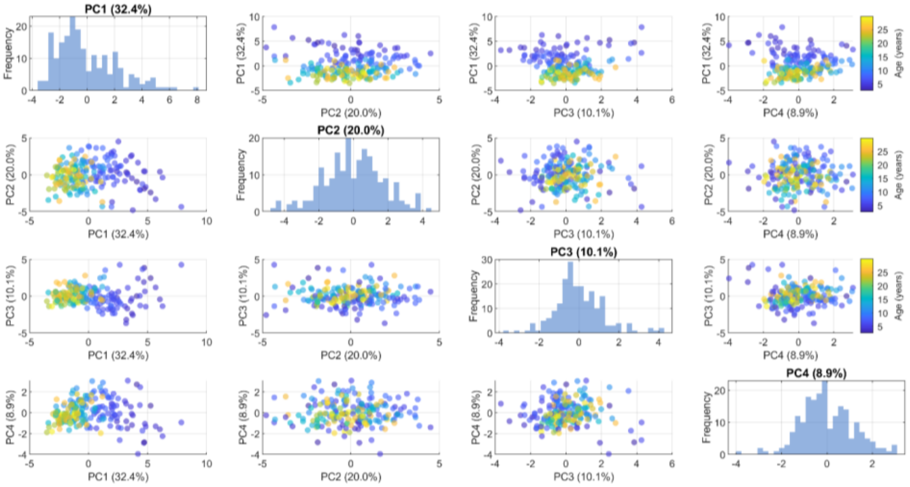


**Figure S1:** Pairwise relationships between principal component (PC) scores derived from the global principal component analysis. Scatter plots display the distribution of observations across PC1-PC4, with color indicating age (years). Histograms along the diagonal represent the distribution of scores for each principal component. Percentages in parentheses denote the proportion of total variance explained by each component.
